# Pore-like diffusion barriers in murine cardiac myocytes

**DOI:** 10.1101/2023.01.02.522313

**Authors:** Christine Deisl, Jay A. Chung, Donald W. Hilgemann

**Affiliations:** Department of Physiology, Southwestern Medical Center, Dallas, TX 75235-9040 USA; Laboratory of Obesity and Aging Research, Cardiovascular Branch, NHLBI, NIH, Bethesda, MD 20892, USA

## Abstract

Using both optical and electrical methods, we document that solute diffusion in the cytoplasm of BL6 murine cardiac myocytes becomes restricted >30-fold as molecular weight increases from 30 to 2000, roughly as expected for pores with dimensions of cardiac porin channels. The Bodipy-FL ATP analogue diffuses ∼50-fold slower in BL6 cardiac cytoplasm than in free water. From several fluorophores analyzed, our estimates of bound fluorophore fractions range from 0.1 for a 2 kD FITC-labeled polyethylene glycol to 0.93 for sulforhodamine. We estimate that diffusion coefficients of unbound fluorophores range from 0.5 to 8 x 10^-7^ cm^2^/s. Analysis of Na/K pump and veratridine-modified Na channel currents confirms that Na diffusion is nearly unrestricted (time constant for equilibration with the pipette tip, ∼20 s). Using three different approaches, we estimate that ATP diffuses 8 to 10-times slower in the cytoplasm of BL6 myocytes than in free water. To address whether restrictions are caused more by cytoplasmic protein or membrane networks, we verified first that a protein gel, 10 gram% gelatin, restricts solute diffusion with strong dependence on molecular weight. Solute diffusion in membrane-extracted cardiac myofilaments, confined laterally by suction into large-diameter pipette tips, is however less restricted than in intact myocytes. Notably, myofilaments from equivalently extracted skeletal (diaphragm) myocytes restrict diffusion less than cardiac myofilaments. Solute diffusion in myocytes with sarcolemma permeabilized by β-escin (80 µM) is similarly restricted as in intact myocytes. Diffusion restriction in cardiac myocytes is strain-dependent, being about two-fold greater in BL6 myocytes than in myocytes with a CD1/J6/129svJ background. Furthermore, diffusion is 2.5-fold more restricted in CD1/J6/129svJ myocytes lacking the mitochondrial porin, Vdac1, than in WT CD1/J6/129svJ myocytes. We conclude that both myofilaments and mitochondria networks restrict diffusion in cardiac myocytes. As a result, long-range solute diffusion may preferentially occur via passage through porin channels and intramembrane mitochondrial spaces, where diffusion is less restricted than in myofilament spaces.

## Introduction

The evolution of cellular life was shaped profoundly by the physical properties of diffusion in aqueous solutions (Gallet et al., 2017). Diffusion limits the dimensions of cells and determines how cells can be organized into viable three-dimensional tissues. The circulatory system can be viewed as an evolutionary response to the limits of diffusion (Popel, 1989). Nevertheless, the magnitudes of ion and solute gradients during many physiological processes remain poorly established, and significant questions remain as to how local diffusion is influenced by cellular protein matrices and membrane networks (Wheatley, 2003; Sanabria et al., 2007). For skeletal muscle, one of the most influential analyses appeared in 1969 (Kushmerick and Podolsky, 1969). As expected from the fact that Ca is heavily buffered by Ca-binding proteins, Ca diffusivity is reduced 50-fold in muscle versus free water. In contrast, monovalent ions and ATP were shown to diffuse only about two-fold less well than in free water. While these impressions have by-and-large been supported for skeletal muscle, several studies highlighted next suggest that solute diffusion is more restricted in cardiac myocytes and requires further attention.

Diffusion of sodium ions in cardiac myocytes has been controversial for decades, a key proposal being that cytoplasmic Na diffusion is selectively restricted in subsarcolemmal spaces (Carmeliet, 1992), such that large changes of local Na concentrations occur during Na transport (Verdonck et al., 2004). Although subsarcolemmal Na gradients are now questioned (Swietach et al., 2015; Garcia et al., 2016; Lu and Hilgemann, 2017), cardiac Na channels (NaV’s) and Na/K pumps are still proposed to generate local Na gradients that require many seconds, even minutes, to dissipate (Skogestad et al., 2019). Therefore, we reexamined this issue and verify here that Na diffusion in cardiac myocytes is effectively unrestricted.

The diffusion of organic solutes within protein gels, as well as other polymers, is highly complex (Masaro, 1999; Sanabria et al., 2007). Already in 1930 it was described that organic solutes encounter diffusion barriers in fine-grained gelatin with size exclusion behavior expected for 1 to 2 nanometer pores (Friedman, 1930). For muscle, the diffusion of nucleotides naturally received more attention than other solutes, given the roles of nucleotides in energy metabolism (Saks et al., 1994; Jepihhina et al., 2011) and cell signaling (Jones, 1986; Weiss and Lamp, 1987, 1989). Von Wilhelm Hasselbach reported in 1952 that the diffusion coefficient of ATP in skeletal muscle was ∼3·10^-8^ cm^2^/s (Hasselbach, 1952), 100 times smaller than in free water (3 to 7 x 10^-6^ cm^2^/s, in the presence and absence of monovalent ions, respectively) (Bowen and Martin, 1964). The coefficient given by Hasselback was later questioned and corrected for glycerol-treated muscle to a value of 2×10^-6^ cm^2^/s or about one-half the free diffusion of ATP in water (Bowen and Martin, 1963). This corrected value is essentially the same as suggested later for intact skeletal muscle (Kushmerick and Podolsky, 1969). Baylor and Hollingworth used a coefficient of 1.4 x 10^-6^ cm^2^/s in their 1998 model describing how Ca buffering by ATP can facilitate cytoplasmic Ca diffusion in frog skeletal muscle (Baylor and Hollingworth, 1998). The diffusion coefficient of a non-hydrolyzable ATP analogue in a muscle extract was found to be somewhat smaller, namely 10^-6^ cm^2^/s (Sidell and Hazel, 1987). Importantly, ^31^P NMR studies of ATP in skeletal muscle (Hubley et al., 1995; de Graaf et al., 2000) are overall consistent with these results and suggestions. Notably, the latter study (de Graaf et al., 2000) employed fibers with a low mitochondrial content and showed that ‘instantaneous’ ATP diffusion occurs with no restriction at all, while diffusion over long distances becomes restricted by about 60% with apparent boundaries occurring at distances of 15 to 22 µm. As described next and verified in our experimental results, this pattern of long-range diffusion restriction appears to be more pronounced in cardiac myocytes than in skeletal muscle.

The existence of cytoplasmic free ATP gradients in a variety of cell types has been indirectly supported by multiple experimental methods. Functionally important cytoplasmic ATP gradients were indicated by ‘cryomicrodissection’ studies in oocytes (Miller and Horowitz, 1986). Relevant to membrane transport, membrane cytoskeleton was proposed to define pools of ATP in red blood cells that are functionally distinct from bulk cytoplasmic ATP (Chu et al., 2012). In other words, protein networks, as well as membrane networks, are proposed to restrict diffusion of organic solutes to a physiologically important extent.

Multiple studies suggest that diffusion of organic solutes is substantially restricted in cardiac myocytes: The diffusion coefficient of a fluorescent ATP analogue was determined in rat cardiac myocytes to be as small as suggested for ATP itself by Hasselbach, namely 0.7-4 x 10^-8^ cm^2^/s (Illaste et al., 2012). This result was interpreted in terms of lattice-like diffusion obstructions in cardiac myocytes. Specifically, diffusion barriers would occur at ∼1 μm distances with barrier pores making up only 0.3% of the barrier surfaces. The barriers were suggested to be membranous in nature, with mitochondria, T-tubules and sarcoplasmic reticulum potentially all taking part. Using sarcolemma-permeabilized cardiac myocytes, this group also showed that cytoplasmic ADP diffusion is restricted 6- to 10-fold (Simson et al., 2016). Similar ATP derivatives, other fluorophores and absorbance dyes appear to diffuse in a substantially restricted manner in cardiac muscle, and to a lesser restricted extent in skeletal muscle. Free (unbound) Fura-2 diffuses with a diffusion coefficient of 0.52 x 10 ^-6^ cm^2^/s in skeletal muscle (Pape et al., 1993), just 10-times slower than it diffuses in free water (Timmerman and Ashley, 1986). The absorbance dye, arsenazo III has a free diffusion coefficient of 0.8 x 10^-6^ cm^2^/s in skeletal muscle with about 90% of the dye being bound. Using a photobleach method, the *apparent* diffusion coefficients for free Fura-2 and Indo-1 Ca dyes in guinea pig cardiac myocytes were determined to be 1.6 and 3.2 x 10^-7^ cm^2^/s, respectively (Blatter and Wier, 1990). Assuming that these dyes diffuse at about 10^-6^ cm^2^/s in free water, these authors concluded that the fluorophores diffuse only 3 to 6 fold slower in myoplasm than in free water. In fact, the diffusion coefficient of Fura-2 in water had been determined to be 4.7 x 10^-6^ cm^2^/s (Timmerman and Ashley, 1986), indicating that dye diffusion is 15 to 30 times slower in guinea pig cardiac myocytes than in free water.

The existence of cyclic nucleotide gradients in many cell types has been inferred for decades on the basis of both functional and optical studies (Hayes and Brunton, 1982; Jurevicius and Fischmeister, 1996; Agarwal et al., 2016). Although buffering of cAMP by its binding sites and hydrolysis of cAMP by phosphodiesterases likely support these gradients, additional factors are also thought to be involved (Agarwal et al., 2016). It is proposed for cardiac myocytes that diffusion of all molecules larger than ions becomes restricted about 10-fold as a result of ‘cytoplasmic tortuosity’ (Richards et al., 2016). Specifically, these authors found that diffusion coefficients of multiple fluorophores in cardiac myocytes were ∼4 x 10^-7^ cm^2^/s, and they proposed that intracellular membranes, in particular mitochondrial membranes, would likely account for diffusion restriction. Notably, this same group found with similar optical methods that Na diffusion was nearly unrestricted (Swietach et al., 2015). Why ions would experience much less tortuosity than small organic solutes was not explained. Similar to fluorescent ATP analogues, fluorescent cAMP analogues are described to diffuse about 50 times more slowly in the cytoplasm of cardiac myocytes than in free water (Sidell and Hazel, 1987). Also similar to results for ATP analogues, diffusion of cAMP would appear to be restricted in a cell-specific manner. In olfactory cilia, for example, cAMP is described to diffuse in an unrestricted manner (Chen et al., 1999).

With this background, we carried out new experiments to characterize diffusion of multiple commonly employed fluorophores in murine cardiac myocytes, we devised protocols to indirectly characterize diffusion of native solutes in myocytes, and we devised experiments to compare diffusion of monovalent anions of different molecular weights in membrane-extracted and sarcolemma-permeabilized cardiac myocytes. We provide evidence that myofilaments themselves can restrict diffusion, but we also show that the outer mitochondrial membrane importantly restricts diffusion in the cytoplasm of cardiac myocytes. We describe that diffusion restrictions depend on the strain of mice from which myocytes are isolated and that knockout of the mitochondrial porin channel, Vdac1, decreases the apparent diffusion coefficients of ATP and fluorophores by more than two-fold. Given that diffusion restrictions in cardiac myocytes are imposed by both myofilaments and mitochondria, we suggest that the combined restrictions promote cytoplasmic solute diffusion to occur through intermembrane mitochondrial spaces.

## Methods and Principles

### Murine myocytes and patch clamp. The UT Southwestern Medical Center Animal Care and Use Committee approved all animal studies

Cardiac myocytes were isolated from adult (2.5 to 4 month old) mice of either sex and patch clamped as described previously (Lu et al., 2016). Diaphragm myocytes were isolated by gently agitating excised pieces of diaphragm muscle for 30 min in the same solutions employed for cardiac myocyte isolation. Axopatch 1C patch clamp amplifiers were employed using our own software (Wang and Hilgemann, 2008). Unless stated otherwise, solutions had the same compositions as employed previously (Lu et al., 2016; Lu and Hilgemann, 2017). As indicated, N-methyl-D-glucamine (NMG) was used as the substitute for extracellular Na. All experiments were performed at 0 mV. Na/K pump-related experiments were performed at 35°C. Pump currents are activated by exchanging 7 mM extracellular Na for 7 mM extracellular K. The presence of 7 mM Na in the K-free solution is required to suppress pump activation by contaminating K in routinely available chemicals (Lu et al., 2016). Conductance measurements were performed with our own software (Wang and Hilgemann, 2008), employing either sinusoidal or square-wave voltage oscillations of 3 to 10 mV at 0.1 to 0.5 KHz.

Extracellular solution changes were performed by abruptly moving the microscope stage so as to place the myocyte directly in front of 1 of 4 square pipettes (1 mm) with solution steams maintained by gravity-driven flow at velocities of 2 to 5 cm/s. The rapidity of solution changes was quantified from the deactivation of Na/K pump currents upon removing extracellular K. The moment when myocytes crossed the interface between two solutions was detectable as a small current instability, and the subsequent time required for pump current to decay by >50% amounted to 201±8 ms. Since the half-maximal K concentration is less than 0.3 mM, this corresponds to ∼4 half-times of solution exchange. For optical experiments, we patch clamped myocytes close to one myocyte end so that the diffusion path constituted most of the length of the myocytes.

### Transgenic Mice

Knockout of VDAC1 has partial embryonic lethality in the BL6 mouse strain (Weeber et al., 2002). Therefore, VDA1 knockout mice were bred in a CD1/J6/129svJ strain.

### Diffusion models

Simulation and experimental models employed in this study are illustrated in Fig. 1. Diffusion was simulated in one dimension, as shown in Fig. 1A and described subsequently. Experimental models employed are illustrated in Figs. 1B to F. **Patch clamp of intact myocytes.** Fig.1B illustrates patch clamp of murine cardiac myocytes from close to one end to generate a diffusion distance of ∼120 µm. **Membrane-permeabilized myocytes.** As shown in Fig. 1C, membrane-permeabilized myocytes were aspirated into highly polished, large-diameter pipette tips to monitor diffusion through a restricted myofilament. To do so, borosilicate pipette tips were cut and melted to generate bullet-shaped tips with thick (>5μm) terminal walls and openings of 6 to 12 microns (Hilgemann and Lu, 1998). Myocytes selected with a receding appendage, allowing them to be rapidly sucked into the pipette tip in a mechanically stable fashion. Using intact myocytes, the seal resistances achieved with standard recording solutions were minimally 20 MΩ. For some experiments, the myocyte sarcolemma was permeabilized during experiments with the saponin, β-escin (80 μM). To characterize diffusion restrictions conferred by myofilaments themselves, membranes were extensively extracted for 2 to 10 days in solutions consisting of 50% glycerol with 1 mM Triton X-100, 1 mM EGTA, and 2 mM MgATP at 4°C. Additional details are provided subsequently. As illustrated in Fig. 1C, we estimate that the restricted diffusion path amounted to distances of 20 to 30 μm.

**Figure 1.**
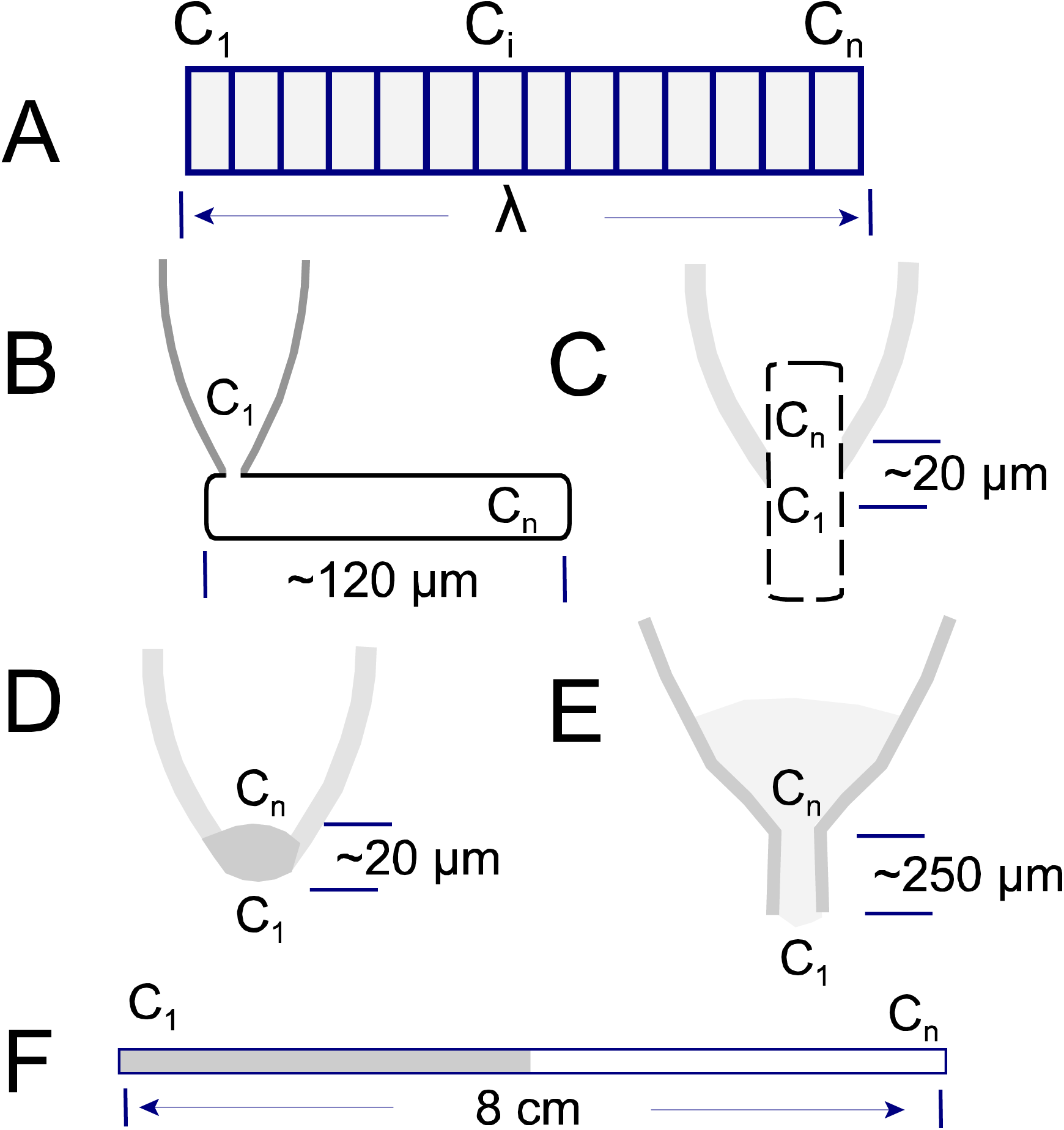
Diffusion models employed in this study. **A.** Simulations were carried out in one dimension with concentrations at the onset (C_1_) being fixed, the concentration at the end point (C_n_) being either free or fixed, depending on the experimental model approximated. Ten to 30 discrete compartments were employed. **B.** Myocytes were patch clamped close to one end, selecting myocytes rangin in length from 110 to 130 µm. For simulatinos, concentrations of solutes within the pipette tip were assumed to be constant. **C.** Permeabilized or detergent/glycerol-extracted myocytes were aspirated rapidly into highly polished pipette tips with inner diameters of ∼12 µm. The restricted region, limiting conductance of the patch pipette, was estimated to be 15 to 30 µm long. **D and E.** Gelatins and/or viscous polymer solutions were aspirated into pipette tips, and the limiting conductance was monitored during and after rapidly switching the pipette tip between solution streams of different compostion. Pipettes were employed with restrictions occuring over about 20 microns (D) up to 300 microns (E). **F.** Eight centimeter long glass pipettes with a 1 mm inner diameter with carefully filled over 4 cm with one chosen solution, and the corresponding end was sealed with dental wax. Then, the second half of the pipette was filled with a second chosen solution, the corresponding end was also sealed with wax, and fluorescence profiles were aquired at chosen times as described in the text.

### Diffusion through viscous solutions containing macromolecules

Figures 1D to 1F illustrate three means employed to characterize solute diffusion through viscous solutions of macromolecules. Results from these experiments are described in Supplemental Data. First, such mixtures were aspirated into the tips of pipettes similar to those used with myocytes. In addition, as shown in Fig. 4E, we employed 1 mm diameter borosilicate pipettes whose tips were melted on a glass flame, so as to generate nearly linear vestibules with lengths of 0.25 to 0.5 mm and diameters 30 to 60 μm. Micrographs are provided in Sup. Fig. 4. As shown in Fig. 1F, for optical measurements of diffusion over large distances, either in free water or in the presence of macromolecules, borosilicate glass capillaries with inner diameters of 1 mm and lengths of 8 cm were filled by negative pressure over a 4 cm distance with a chosen solution of desired composition. The corresponding end was sealed by dipping into molten dental wax and allowing the wax to harden. Then, the remaining half of the pipette was filled via a fine polyethylene tubing (∼0.1 mm outer diameter) taking care to avoid solution mixing. The distal end of the tube was then also sealed with molten dental wax. Finally, to measure the conductance of solutions in relation to fixed dimension, we employed 15 cm long lengths of Tygon® tubing with an inner diameter of 0.5 mm (not shown). Solutions were aspirated into the tubes by negative pressure, and the ends were placed into chambers containing 30 mM KCl solutions with silver/silver chloride electrodes for conductance measurements via patch clamp.

When gelatins were employed, they were prepared as an equal mix of Type A and Type B gelatins (Sigma-Aldrich). The gelatins were cleaned by allowing them to set first in a 10 cm diameter beaker at a thickness of ∼0.5 cm. After setting, gelatins were soaked with a chosen solution and tilt-rocked for 2 to 3 days. The gelatin was then re-liquified by heating and was allowed to set for 10 min in pipettes before filling the empty pipette region with gelatin-free solution. Fluorophores were added to either the gelatin solution or the gelatin-free solution, or both, as indicated in relevant Supplemental Data. After intervals of an hour to 7 days, the tubes were imaged with a Biorad Chemi-DOC MP imaging system to assess diffusion of fluorophores or fluorophore ligands.

### Concentration-conductance relations relevant to this study

Figure 2A shows a log-log plot of the concentration-conductance relationship obtained for KCl in 15 cm long, 0.5 mm diameter tubes. Over three log units, from 40 µM to 200 mM, the concentration-conductance relation appears linear. When plotted with a linear concentration scale, however (Fig. 2B), the well-known non-linearity of this relationship is apparent. The results in Fig. 2B are fitted by assuming that the conductance (G) reflects a simple dissociation reaction of KCl with the dissociation coefficient, K_d_,

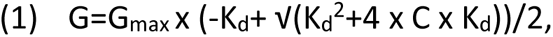

**Figure 2.**
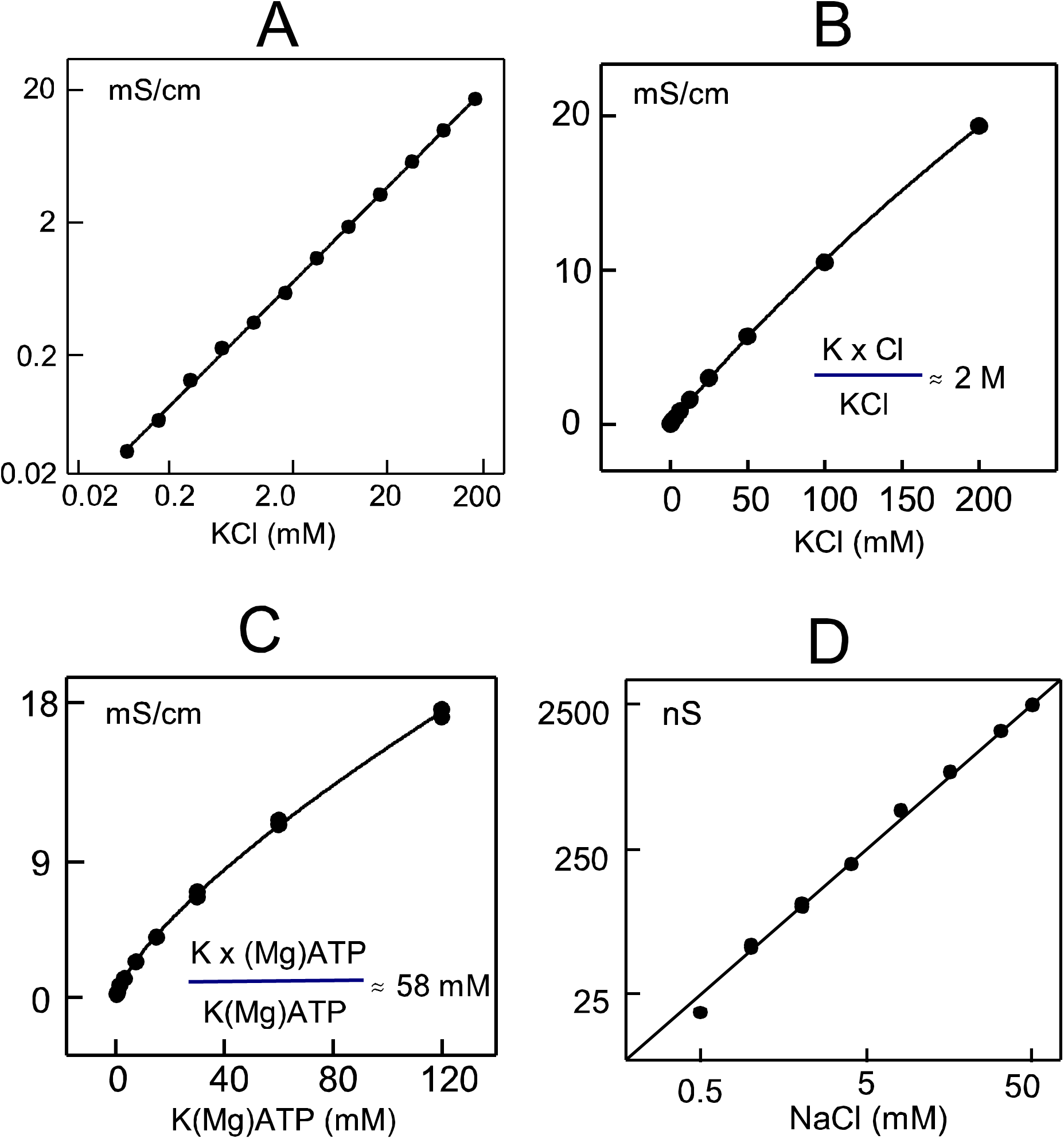
Concentration-conductance relations of ionic solutes employed in this study. **A-C.** Conductivity of solutions aspirated into 15 cm long Tygon® tubes with an inner diameter of 0.5 mm. **A.** A log-log plot of the conductivity of KCl solutions appears linear over three log units. **B.** Non-linearity of the KCl concentration-conductance relation is apparent in a linear plot. The relation is reasonably described by a KCl dissociation constant (k_d_) of 2 M, fitting results to the equilibrium solution of the reaction (K x Cl)/KCL = k_d_. With C as the concentration of total KCl, G (mS/cm)= G_max_ x (-k_d_+√(k_d_^2^+4C))/2. **C.** Concentration-conductance relation of K(Mg)ATP solutions, prepared by dilution of a 120 mM MgATP solution, titrated to pH 7.0 with KOH. Non-linearity over the concentration range from 0 to 120 mM is well described by a K(Mg)ATP dissociation constant of 58 mM. **D.** Concentration-conductance relation of NaCl solutions from 0.5 to 50 mM, determined by aspirating solutions into a patch pipette tip with a large diameter. Nonlinearity (not apparent in the log-log plot) is similar to that of KCL solutions.

where C is the concentration of salt added to solution. A dissociation constant of about 2 M accurately predicts that 8% of KCl is undissociated in a 200 mM KCl solution. As needed to characterize ATP diffusion via conductance changes, Fig. 2C shows the equivalent titration of K(Mg)ATP, diluted from a 120 mM stock solution of MgATP that was set to pH 70 with KOH. The relationship is substantially more non-linear than for KCl, and the best fit to equation #1 has a dissociation constant for K(Mg)ATP of 58 mM. For measurement of NaCl diffusion, Fig. 2D underscores via a log-log plot that nonlinearity of the NaCl concentration-conductance relation, determined in this case by aspirating solution the tip of a patch pipette, is similar to that of KCl. Both experimental (Artemov, 2015; Kamceva, 2018; Widodo, 2018; Lee, 2020) and theoretical (Lee, 2020) studies show that non-linearities o the relationship amount to less than 10% under conditions of our experiments. In conclusion, the nonlinearity, predicted by Kohlrausch (Lee, 2020), i.e. the dependence of molar conductivity (Λ_m_) on the limiting molar conductivity (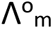) as a function of concentration (c, Λm=Λom-K·c^1/2^), can be ignored for KCl and NaCl, but not for K(Mg)ATP, in our experiments.

### Simulations of diffusion in one dimension

For all simulations of diffusion, 10 to 30 discrete compartments were assumed (see Fig. 1A) with diffusion between compartments determined by a diffusion coefficient, D_x_ with no dependence on concentration:

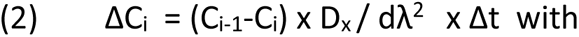

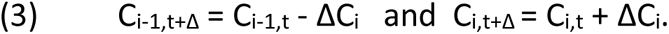

Simulations were performed with dλ in cm, D_x_ in cm^2^/s and concentrations in mM. As outlined previously (Lu and Hilgemann, 2017), we estimate the cytoplasmic mixing volume of murine cardiac myocytes be ∼12 pL, which corresponds to ∼30% of their geometrical volume (∼36 pL). In all simulations, the concentration of the 1^st^ compartment was fixed and changed at given times to simulate addition or removal of solutes from the pipette tip. For cardiac myocytes (Fig.1B), the diffusion constant at the C_1,2_ interface was reduced to mimic an input resistance that was 3 to 4 fold greater than the total longitudinal myocyte resistance with 120 mM KCl. For myocytes, the nth compartment communed only with the n-1 compartment, while for other models the concentration of the nth compartment was fixed at the solute concentration included initially in the pipette. For simulations of diffusion in long (8 cm) pipettes, the initial condition was that the two halves of the pipette contained two different solute concentrations.

All simulations enforce electroneutrality and implement Kohlraush’s principle that solution conductivity is *proportional* to the sum of the products of all free ion concentrations with their respective diffusion coefficients. For routine purposes, we did not calculate absolute conductances. As shown in FIg. 3, the coupled diffusion of a single monovalent anion/cation pair (X^+^Y^-^) is accurately described by the coupled diffusion coefficient, D_XY_, where

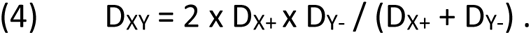

To demonstrate this and to allow simulation of all monovalent ions commonly employed in experiments, Na, Cl, K and one anion (A), the diffusion equation for each ion is modified by an electrochemical factor, Ke, corresponding to the local electrical field as e^0.5*z*ΔΨxF/RT^. Assuming that A is monovalent, then Ke is calculated as follows:

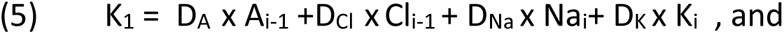

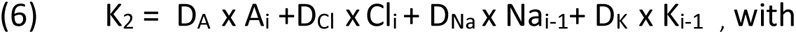

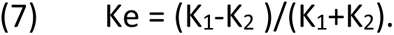

**Figure 3.**
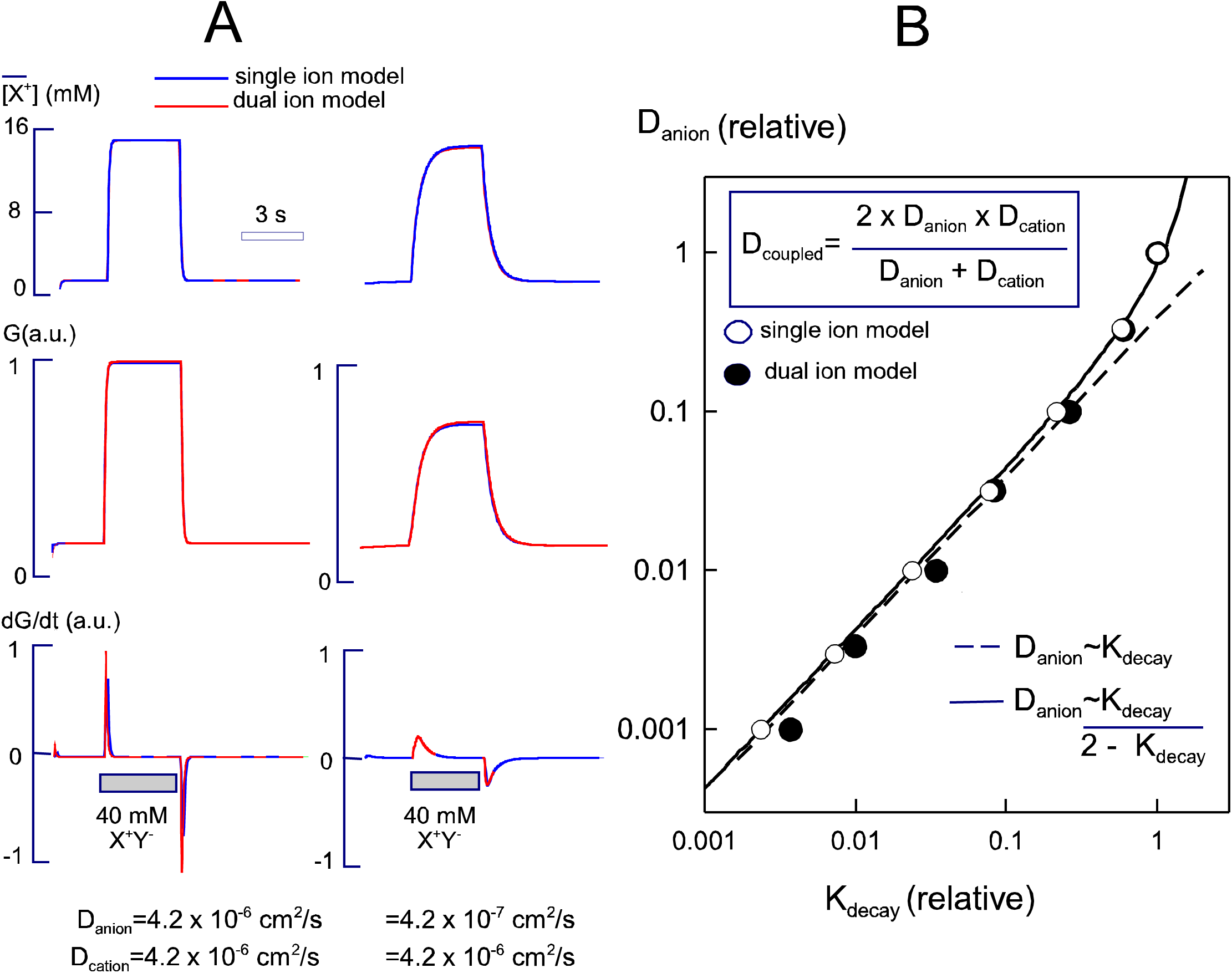
Simulated average ion concentrations and conductance of a column 10 µm long, open at both ends, with diffusion taking place equivalently over an additional 5 µm at both ends, outside of the restricted zone. **A. Left.** Results for an ion pair with equal anion and cation diffusion coefficents. Initially, the concentration is 2 mM everywhere. As indicated the concentration was increased to 40 mM at the front of the column for 3 s and then decreased again to 2 mM. **Right.** Equivlanet results for for the case that the anion diffusion coefficient was 10-fold less than the cation diffusion coefficient. Results in red are a model in which diffusion of the anion and cation were simulated separately, results in blue are for a model assuming a diffusion coefficient for the ion pair as given in panel B. **B.** Plot of the rate constants (X-axis), determined from single exponential fits of the decaying conductance, against the relative anion diffusion coeffienct (Y-axis), which was varied over 1000 fold. The predicted coupled diffusion constant accurately describes the relationship over the entire three log unit range.

Employing these equations for one anion and one cation, it is shown in Fig. 3A that results employing a coupled diffusion coefficient (3) and the equation set (5 to 7) generate essentially the same results, here for 40 mM of an ion pair added to and removed from a base solution with 1 mM of the same ion pair. Diffusion was simulated over a distance of 120 μm through a open-ended column, as drawn in Fig. 1A. The left and right panels show results for identical anion and cation diffusion coefficients and for a 10-times lower anion coefficient, respectively. From top to bottom, the plots give the average ion concentration, the relative conductance of the diffusion column, and the first derivative of the conductance. Also plotted but not visible, single exponential functions can be fit accurately to the rising and falling conductance curves over 80 to 90% of their time course (red versus blue curves). As shown in Fig. 2B, the rate constants determined in this manner (X axis) can be used to determine the anion diffusion coefficient (Y axis). To do so, equation 3 is rewritten to calculate the anion diffusion coefficient from the rate constants of conductance decay, which in turn can be demonstrated to be proportional to the coupled diffusion constant.

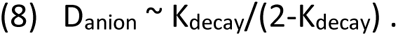

As the anion diffusion coefficient is decreased, the decay rate decreases as indicated in Fig. 2B. The coupled coefficient decreases at first approximately with the square root of the diffusion coefficient. Then, as the anion diffusion coefficient decreases more than a factor of 10, the decay constants decrease linearly with the diffusion coefficient.

To evaluate Na diffusion and accumulation in myocytes, simulations were extended to include a surface membrane Na current. To evaluate additionally MgATP diffusion into myocytes from the pipette tip, simulations were developed for four ions. For a patch clamped myocyte at 0 mV, the Na current was simply assumed to be proportional to the Na concentration difference across the plasmalemma. In the absence of extracellular Na, the current is calculated in pA (Ina_i_=Na_i_ x G_Na_), and the electrochemical factor, Ke, is modified to include the cytoplasmic ionic flux (I_i_) as Ke’. The modified factor then reflects the Na current occurring within each compartment plus the sum of Na currents occurring distal to that compartment (∑ Ina_i_ to Ina_n_). The charge flux, calculated in millimole/L/s, is then the local current in pA divided by 100 and the compartment volume in pL:

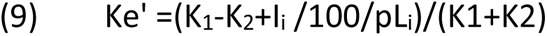

To reconstruct experimental results, four ion concentrations are then simulated as follows where A is an anion with diffusion coefficient, D_A_, Cl is chloride, Na is sodium and K is potassium:

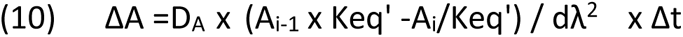

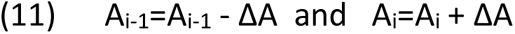

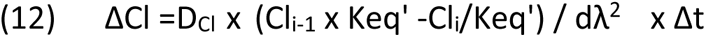

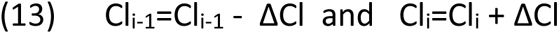

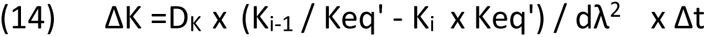

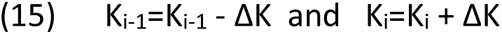

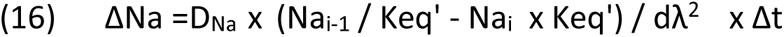

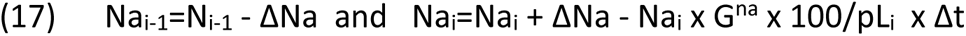

MgATP is often assumed to carry about 2 negative charges at pH 7. Without Mg, the 4^th^ proton site of ATP is ∼1/2 dissociated at pH 7.2 (Stockbridge and Wolfenden, 2009). It has been our routine observation, however, that titration of high-grade MgATP (Sigma-Aldrich) yields pH values higher than 7.0 with addition of 1.2 moles of alkali per mole of MgATP. This is consistent with the 4^th^ protonation site remaining largely undissociated at pH 7.2. ^31^P NMR analysis of MgATP solutions with extended pH titrations supports the conclusion that valence of MgATP at pH 7.0 is about −1.2 (Song, 2008). ^31^P NMR also indicates that ATP dimers, as well as monovalent cation ATP complexes, are likely physiologically significant ATP complexes in cells (Glonek, 1992). All results presented in this article simulate MgATP as a monovalent anion, but results were not significantly different when routines were developed to simulate ATP valences over a range of −1.2 to −2.0.

To simulate diffusion through pipette tips, the total tip conductance is the reciprocal of the total pipette tip resistance. For our purposes, it was adequate to employ relative conductivities, corresponding to the sum of the local ion concentrations, multiplied by their respective diffusion coefficients. Therefore, for simulations of Na, Cl, K and one additional anion,

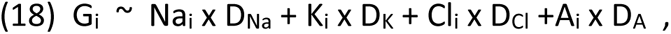

with the total pipette tip conductance, G_total_, being inversely proportional to the sum of the compartment resistances in series:

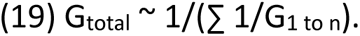

For simulation of pipette tips with long diffusion paths, the pipette dimensions were reconstructed digitally and used to simulate diffusion in the tip through linear compartments with increasing diameters at increasing distances from the tip opening. For simulations of pipette tips with aspirated myocytes, we assumed that the limiting pipette tip conductance arose from a restriction 10 to 30 μm long with diffusion taking place over an additional 5 to 15 μm with similar diffusion characteristics. All simulations were carried out with error checking and step-size adjustment to ensure accuracy within line-width of plotted results.

Figures 4 illustrates one further simulation result that is important for this study, namely an analysis of multiple kinetic components when three ions are present in experiments. In this example, a 20 µm diffusion path is assumed to be open at both ends (approximating Fig. 1D) with 100 mM KCl being present initially on both sides of the confined path. In the hypothetical experiment, 50 mM K(Mg)ATP is applied to one side in addition to 100 mM KCl for 3 s. The diffusion coefficient of the ‘monovalent’ (Mg)ATP anion is varied by 16-fold. For results in Fig. 4A, the diffusion coefficient of ATP is 0.8 x 10-5 cm^2^/s while that of both K and Cl is 2 x 10^-5^ cm^2^/s. The rising and falling conductance curves have fast and slow components, the fast component corresponding to the coupled diffusion constant of KCl and the slow component corresponding to the coupled diffusion coefficient of K(Mg)ATP. In brief, the fast component corresponds to an initial diffusion of KCl into the confined space, as K diffuses ahead of K(Mg)ATP into and out of the confined space. The slow component corresponds to exchange of excess Cl in the confined space for (Mg)ATP, followed by coupled diffusion of K(Mg)ATP. The red curves in Fig. 4A are the best fits of double exponential functions to the simulated conductance curve. The recovered time constants are in good agreement for the rising and falling phases. Fig. 4B plots results of simulations in which the anion (MgATP) diffusion coefficient was decreased by 12-fold from that of Cl. The slow exponential time constant, relative to the time constant with all diffusion coefficients being equal, is plotted on the Y-axis against the assumed diffusion coefficient of MgATP on the X-axis, relative to that of Cl. The slow exponential phase of both the rising and falling conductance curves represent accurately the ratio of the diffusion coefficient of the second anion (i.e. MgATP) to that of KCl (i.e. the time constant of the fast exponential in the two-exponential fit). In Results, these same principles will be exploited to analyze coupled diffusion of Na with Cl, aspartate, MgATP, and an anionic polyethylene glycol.

**Figure 4.**
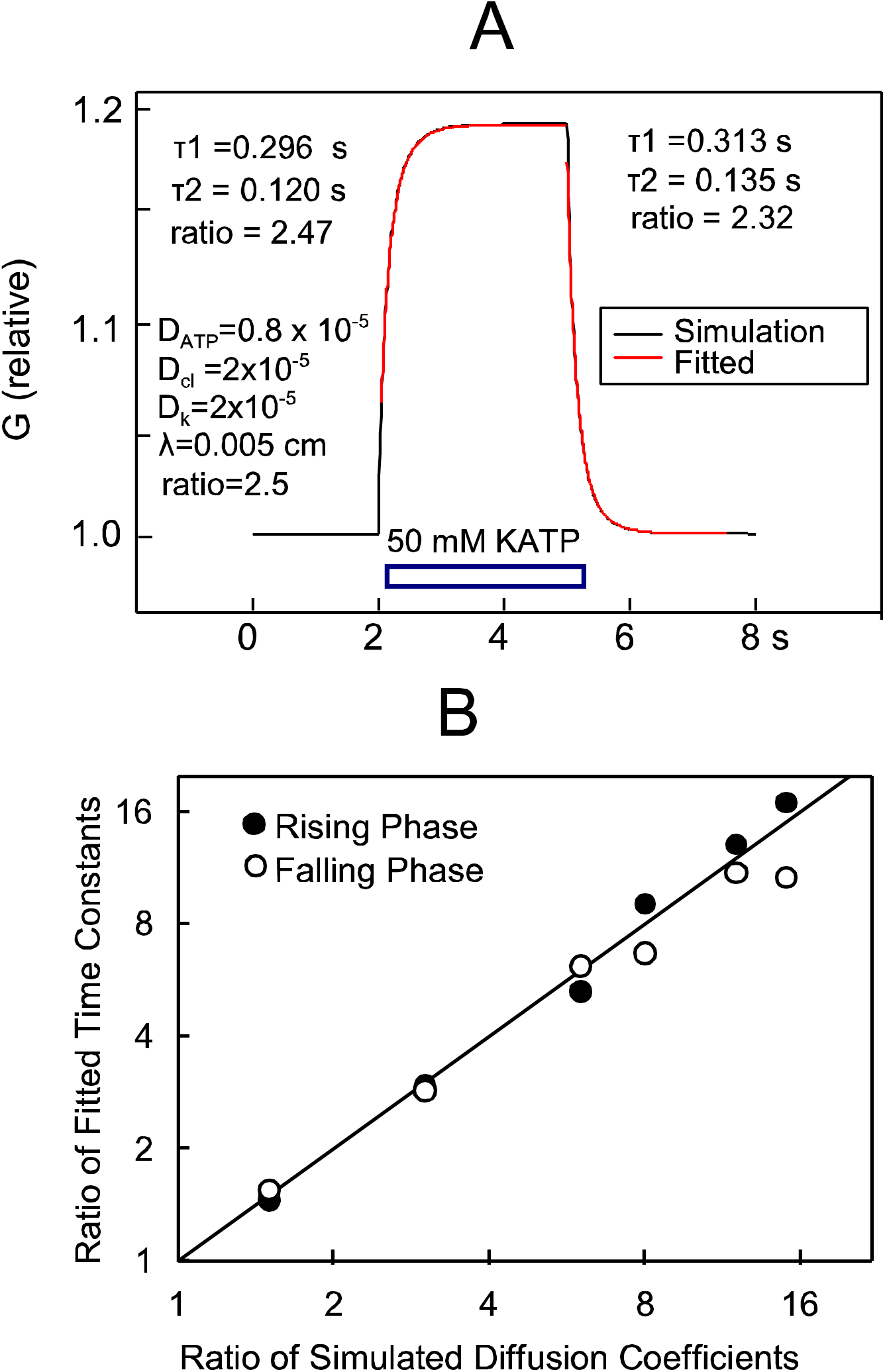
Determination of relative diffusion coefficeints when three ions are present in simulations equivalent to those of Figure 2. A. Initially 100 mM KCl is assumed to be present everywhere, whereby K and Cl have identical diffusion coefficients. As indicated, 50 mM K(Mg)ATP is assumed to be added to the solution in front of the column, whereby the diffusion coefficient of (Mg)ATP is assumed to be 2.5-fold less than that of K and Cl. The results give rise to double exponential rising and falling conductances, whereby the fitted time constants occur in ratios of 2.47 and 2.32, respectively. B. The ratios of time constants fitted to simulations in which the diffusion coefficient of ATP was varied over a 10-fold range with respect to those of K and Cl. The fitted exponentials accurately recover the relative diffusion constants of (Mg)ATP and KCl over the entire concentration range.

### Materials

Unless stated otherwise, chemicals were from Sigma-Aldrich (St. Louis, MO) and were the highest grade available. Unbranched FITC-labelled polyethylene glycols (PEGs) were purchased from Creative PEGWorks (Chapel Hill, NC, 27516). Except for carboxyfluorescein and sulforhodamine, other fluorophores were from Invitrogen. Luciferase was from Creative Biomart (Shirley, NY). Purified GFP was a generous gift of Michael Rosen (UTSouthwestern). VDAC1 knockout mice were bred in a C57BL6 background (Anflous et al., 2001; Kim et al., 2019), and control myocytes for the relevant experiments were of the same sex and background as knockout myocytes.

### Fluorescence imaging of cardiac myocytes

Imaging at near confocal resolution was carried out with an Aurox Clarity laser-free microscopy imaging system (Abingdon, Oxfordshire, OX14 3DB, UK) employing a Nikon Eclipse TE 2000-S inverted microscope. A 40x water immersion lens was employed. All solutions were filtered with 0.22 μm syringe filters to remove aggregates. The times given in figures are the times after myocyte opening by suction. Optical experiments were performed at 25°C. To minimize photo-bleaching, images were taking at intervals of 30 to 60 s employing exposure times as short as possible to allow accurate image analysis, usually 0.2 s.

### Statistics

Statistical significance was assessed via Students T-test after determining that results were distributed normally. Error bars in figures represent standard errors

## Results

### Diffusion of fluorescent dyes in free water revisited

As noted in the Introduction, surprisingly different diffusion coefficients have been assumed for common fluorophores in free water. Therefore, we determined again the diffusion coefficients of several commonly employed fluorophores in water in 8 cm long, 1 mm diameter pipettes. The solution contained 150 mM KCl, 1 mM EGTA and 5 mM HEPES, set to pH 7.4 with KOH. As described in methods, pipettes were filled to 4 cm via negative pressure with either dye-containing solution or dye-free solution. Then, the remaining 4 cm of pipette was filled with the complementary dye-free or dye-containing solution, pipettes were incubated in the dark at room temperature for 1h to 72 h and were imaged as described in Methods. Fig. 5A shows the 1 h and 48 h profiles for Bodipy-ATP in gray and black, respectively. Figs. 5B, C and D show the 48 h profiles in black for Fluo3, Mg-Green and Fluo-5N, employed at concentrations of 1 to 10 μM. In addition, each panel shows the relevant simulations for diffusion coefficients of 10^-6^ cm^2^/s and 4.5 x 10^-6^ cm2/s. In all four cases, including 20 similar experiments, the 48h diffusion profile is described well by a diffusion coefficient of ∼4.5 x 10^-6^ cm^2^ /s, and the profile for 10^-6^ cm^2^/s is ∼4-fold too steep, indicating that it underestimates diffusion of fluorophores in water by about 4-fold.

**Figure 5.**
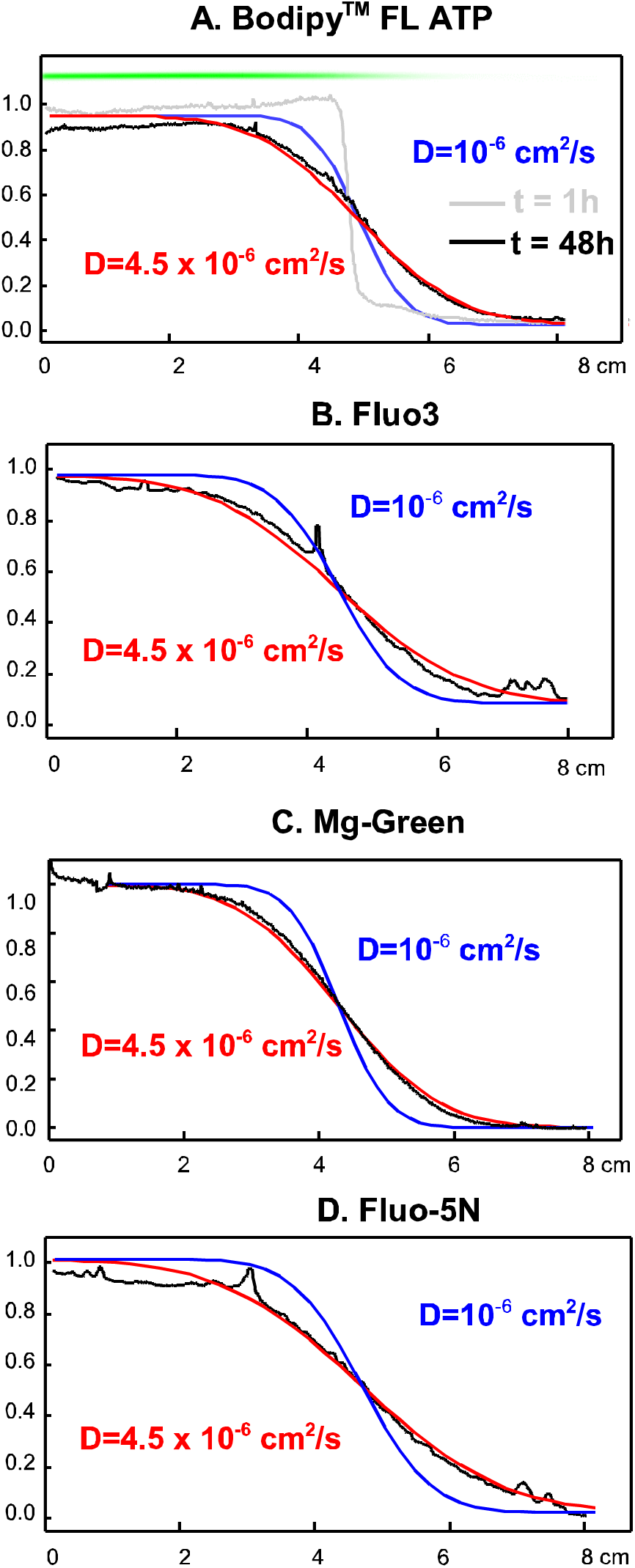
Fluorescence profiles of four fluorophores after initiating diffusion in one-half of 8 cm long glass pipettes with diameters of 1 mm. The solution contained 100 mM KCl, 5 mM HEPES (pH 7.4), and 0.5 mM Mg. For Bodipy FL ATP the fuorescence profile is given at 1 h (grey) and 48 h (black). The fluoresence image for the 48h tube is given above the profiles. For Fluo3, Mg-green, and Fluo-5N, the profiles are given only for 48 h (black). Simulated diffusion profiles for 48 h are given in each panel for a 10^-6^cm^2^/s diffusion coefficient and for a 4.5 x 10^-5^ cm^2^/s diffusion coefficient. In all cases, the larger diffusion coefficient accurately describes the profiles, while the smaller coefficient does not.

### Diffusion of Bodipy-FL-ATP in myocytes

Figure 6 describes the time course with which the fluorescent ATP analogue, Bodipy-FL-ATP, enters a mouse myocyte from a patch pipette and diffuses from one end to the other of the myocyte. Micrographs of the patch pipette tip are shown just after opening the myocyte, as well as at 30 and 60 min time points (A-C). Total fluorescence of the myocyte increases to a plateau with a time constant of 16 min (D). Even after 60 min, line scans reveal longitudinal Bodipy-Fl-ATP gradients in the cytoplasm, decreasing from the pipette tip to the opposite myocyte end (E). Fluorescence of the myocyte cytoplasm protrudes clearly into the pipette tip (see micrograph B). Line scans across the pipette and the width of the myocyte (F) show that the accumulation of dye in extracted cytoplasm versus the pipette tip, itself, amounts to about 30%. Furthermore, fluorescence of the cytoplasm proper does not exceed one-half of the pipette fluorescence, even after 1 h. Accordingly, dye binding to cytoplasmic constituents cannot explain the slow time course with which dye equilibrates with the cytoplasm. To simulate qualitatively the time course of equilibration (G) and the longitudinal gradients after 30 and 60 min (H), a diffusion coefficient of not more than 3 x 10^-8^ cm^2^/s is required. As apparent in Fig. 5H, the simple simulation of diffusion through a column 120 μm in length does not reproduce in detail the longitudinal fluorescence gradients (Fig. 4H). Nevertheless, the fact that fluorescence continued to increase beyond 30 minutes in all experiments is unambiguously consistent with a diffusion process >50-fold slower than in free water, clearly confirming the results of others using other techniques to estimate diffusion coefficients (Illaste et al., 2012).

**Figure 6.**
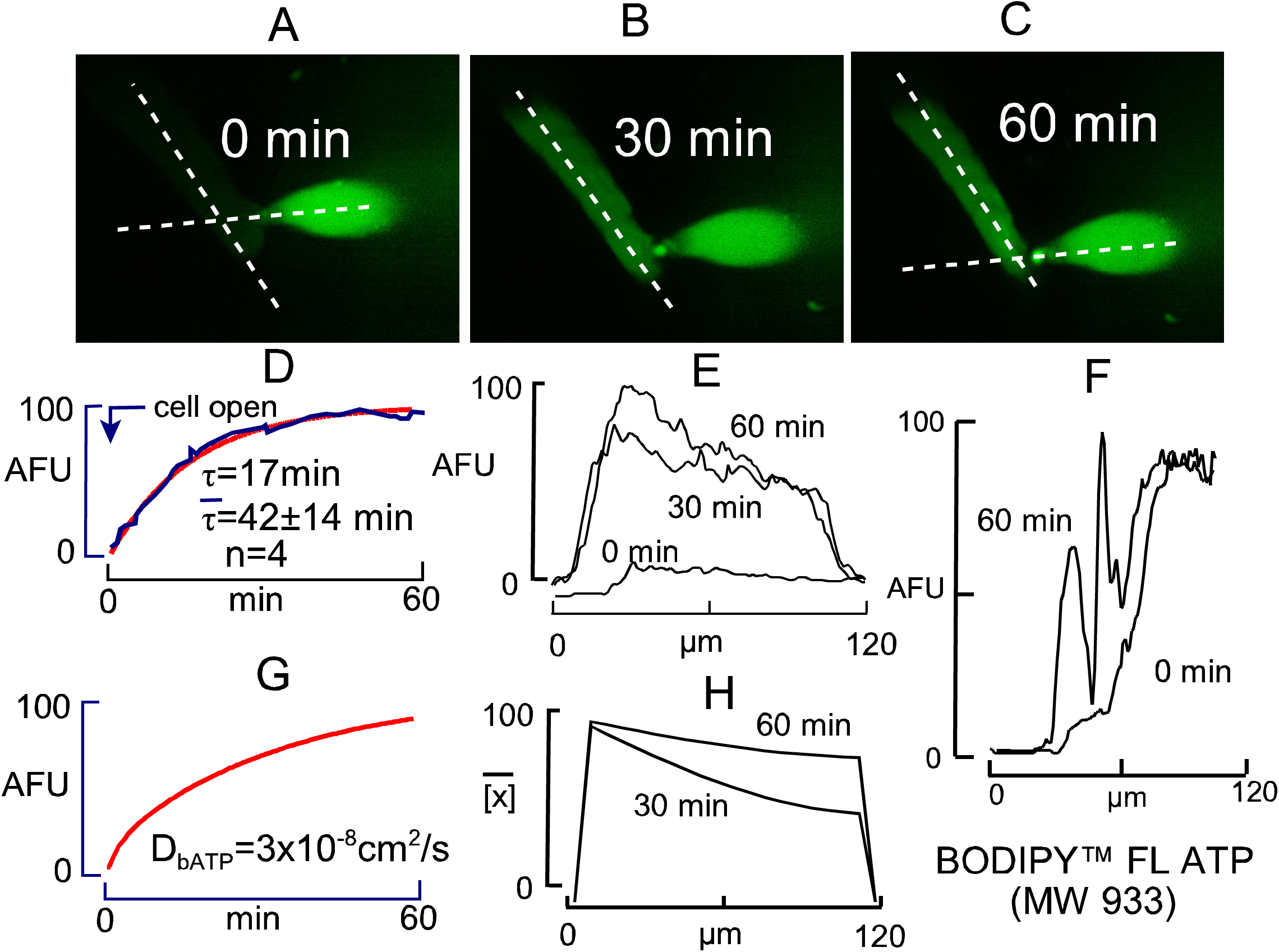
Bodipy^TM^ FL ATP (10 μM) diffusion into a cardiac myocyte from a patch pipette generating a 3 MΩ access resistance. **A-C.** Fluorescence micrographs of the myocyte and patch pipette tip just before opening the myocyte (A), after 30 min (B), and after 60 min (C). **D.** The time course of total myocyte fluorescence change, fitted to an exponential function (time constant, 17 min). The average time constant for 4 experiments was 42 min. **E.** Longitudinal fluorescence line scans along the myocyte at 0, 30, and 60 min. A longitudinal gradient is still evident after 1 h. **F.** Line scan through the pipette tip and across the width of the myocyte, as indicated in A and C, at 0 and 60 min. Note that Bodipy^TM^FL ATP shows a small excess brightness in the pipette orifice where cytoplasm enters the tip. The excess is not more than 12%, however. Therefore, diffusion is not limited by dye binding to myocyte constituents. **G.** Simulation of total Bodipy^TM^ FL ATP equilibration with the myocyte, assuming a myocyte length of 120 microns. From 4 myocytes studied, equilibration was fastest in this myocyte. Therefore, the simulated diffusion coefficient of 3 x 10^-8^ cm^2^/s is the maximal coefficient justified by experimental results. **H.** Simulated Bodipy^TM^ FL ATP gradients.

### Diffusion of all fluorophores is restricted in myocytes

Figure 7 presents equivalent diffusion experiments for 7 fluorophores. Panels A and B present results for sulforhodamine (MW, 559) and carboxyfluorescein (MW, 376). From all fluorophores examined in myocytes, sulforhodomine showed the highest degree of accumulation in cytoplasm with respect to the pipette. This is apparent in the micrograph from the fact that the pipette tip (marked with a diamond) is hardly visible, whereas cytoplasm that is drawn into the pipette tip in line scan #2 is bright. As shown in the middle panel, sulforhodamine equilibrated with the cytoplasm with a very long time constant, on average 28 min (n=5). As evident in line scan #1 in the right panel, the longitudinal gradient was very strong even after 50 min. As apparent in line scan #2, the dye accumulated in the cytoplasm by at least 8-fold with respect to the pipette tip. Obviously, dye binding to cytoplasmic constituents is a strong determinant of sulforhodamine diffusion.

**Figure 7.**
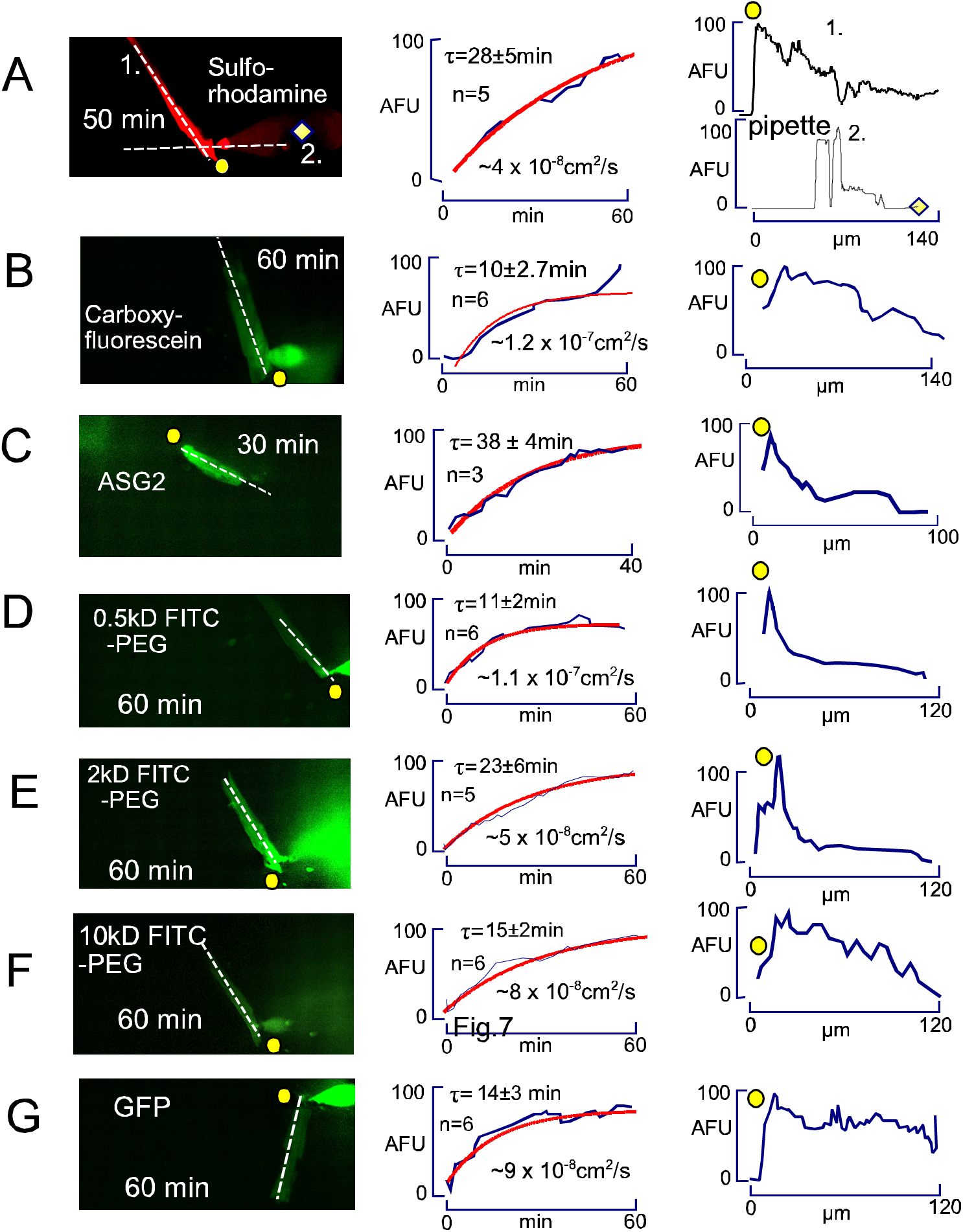
Diffusion of fluorophores into cardiac myocytes during patch clamp with ∼3 MΩ access resistance. Each panel shows the micrograph of a myocyte after approximately 1 h of diffusion (left), the time course of total fluorescence changes during the experiment (middle), and line scans as indicated in the micrograph, using yellow dots to indicate the starting position of the line scan (right). **A.** Sulforhodamine (100 μM) equilibrates with an average time constant of 29 min (middle panel, n=5). It accumulates 8-fold in the cytoplasm (see line scan 2 across the myocyte and into the pipette tip), and it shows a strong longitudinal gradient even after 1 h. **B.** Carboxyfluorescein (100 μM) equilibrates with an average time constant of 10 min (middle panel, n=6), and shows longitudinal gradients even after 1 h. Dye accumulation in cytoplasm versus pipette tip remains very small. **C.** Asante Na-green (ASG-2, 10 μM) also equilibrates with a very long time constant (40 min, middle panel) and shows a longitudinal gradient even after 1 h. Strong fluorescence occurs in myocytes in the absence of Na. **D,E and F.** A 0.5kD, a 2kD, and a 10kD FITC-PEG (100 μM) equilibrate over 1h with average time constants of 11, 23 and 15 min (n=6, 5, and 6, respectively), show no evidence of significant binding, and show clear longitudinal gradients even after 1 h. **G.** Purified GFP (12 μM) accumulates weakly in the cytoplasm with respect to the pipette tip with an average time constant of 14 min (n=6). Fluorescence does not show clear longitudinal gradients.

As shown in Fig. 7B, carboxyfluorescein behaves very differently. The fluorescence of the cytoplasm remains less than that of the pipette tip, indicating that binding to cytoplasmic constituents is weak or negligible. The average time constant for equilibration of dye with the cytoplasm is 10 min (n=6), almost three times less than for sulforhodamine. Notably, the gradient of dye remains substantial after 1 h (see right panel of Fig. 7B). From the rough estimate that D=λ^2^/2/τ, the diffusion coefficient is about 10^-7^ cm^2^/s.

Fig. 7C shows results for the Na-selective dye, Asante Sodium Green-2 (ASG2, MW 1100), equilibrated with a myocyte in the absence of both extracellular and intracellular Na. While this dye exhibits very little fluorescence in a cuvette in the absence of Na, it fluoresces strongly as it diffuses into myocytes in the absence of Na. Clearly, it binds constituents of the cytoplasm and fluoresces as a result. The time constant for dye equilibration is still longer (38 min) than that of sulforhodamine, and the longitudinal gradient remains strong after 1 h. The results demonstrate that ASG2 is NOT a Na-selective dye in the myocyte environment. Since it has negligible fluorescence in the pipette, we cannot estimate the extent to which it binds. However, even if 90% is bound in myocytes, the diffusion constant must be less than 10^-7^ cm^2^/s to account for the time course.

Figures 7D, 7E, and 7F present the diffusion into myocytes of three FITC-labeled PEGs of increasing molecular weight. As shown in Fig. 5D, a 0.5 kD FITC-PEG equilibrates with the cytoplasm with a time constant of 11 min (n=6), more slowly than carboxyfluorescein but more rapidly than sulforhodamine. There was no accumulation in the cytoplasm versus the piptte tip, and the longitudinal gradient remained strong after 1h. Diffusion of a 2 kD PEG (e) was slower by a factor of two (23 min time constant), and a 10 kD PEG (F) diffused similarly to or faster than (time constant, 15 min; n=6) the 2 kD PEG.

Surprisingly, an isolated GFP protein (G) diffused into myocytes with roughly the same time course as the 2 kD PEG (see panel F). Further of note, the GFP fluorescence did not show strong gradients during the 1 h time course examined, and the magnitude of the cytoplasmic fluorescence remained >7-fold less than the GFP fluorescence in the pipette tip. Notably, a similar discrepancy in the diffusion of fluorescent probes with respect to molecular weight in cardiac myocytes was reported previously (Illaste et al., 2012). We speculate in the Discussion that this diffusion process may reflect facilitated transport involving protein-protein interactions. We document in Sup. Fig. S1 that 2 kD PEGs with an added positive amino group or an added negative COOH group did not diffuse differently from the neutral PEG. We also document that an AlexaFluor488-labelled albumin diffused similarly to GFP and that a 4400 MW TRITC-labeled dextran equilibrated similarly to 10K PEG.

Table 1 presents our analysis of the diffusion of 8 fluorophores into murine cardiac myocytes, allowing estimation of the diffusion coefficients for each unbound fluorophore. First, we measured the ratio of fluorescence in the myoplasm next to the pipette tip to that of the pipette at a time point near the end of the experiment. From this ratio, given in column 1, we calculated the fraction of fluorophore likely to be bound, assuming that the cytoplasmic mixing volume is just one-third of the myocyte volume, as determined for monovalent ions (Lu and Hilgemann, 2017). Accordingly, a fluorescence ratio of 0.33 would indicate negligible fluorophore binding to cytoplasmic constituents with the fraction of bound fluorophore (F_bound_) in the cytoplasm estimated as:

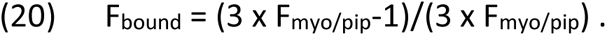

**Table 1.**
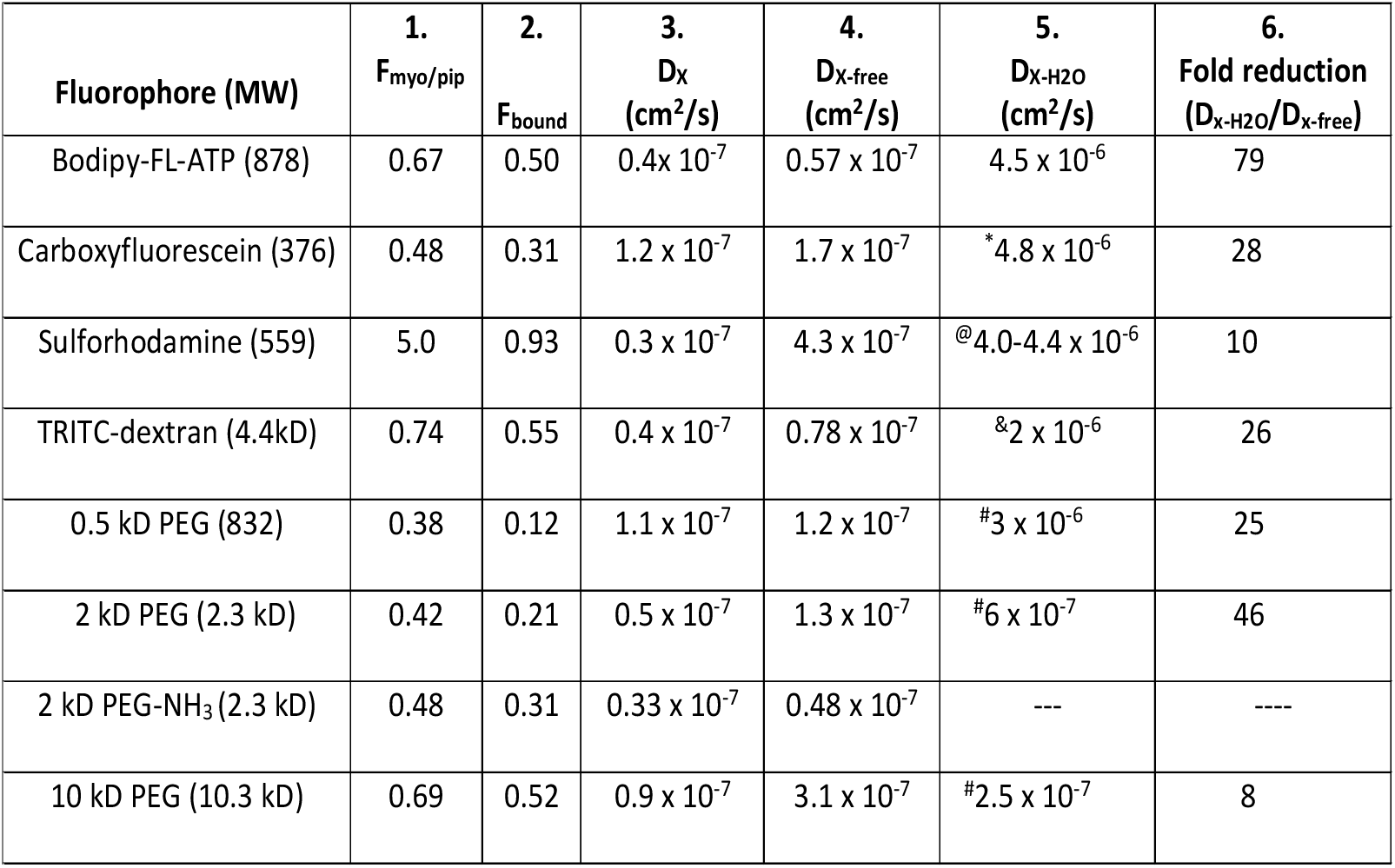
Analysis of fluorophore diffusion in murine myocytes. From left to right: **Column 1.** The ratio of fluorescence of the pipette solution to the fluorescence of myoplasm in close proximity after 1h (F_myo/pip_). Values for BSA and GFP were 0.07 and 0.15, respectively, indicating that they are excluded from most of the myoplasmic space. **Column 2.** Estimated fractions of fluorophores bound by cytoplasm. Assuming a free cytoplasmic space that is one-third of the total myocyte volume (Lu et al., 2016), the fraction of fluorophore bound (F_bound_) is (F_myo/pip_·3 +1)/F_myo/p_·3. **Column 3.** Estimated diffusion coefficients for total fluorophore, estimated as λ^2^/(2·τ) with λ=0.012 cm. **Column 4.** Diffusion coefficients for unbound fluorophore, estimated as D_x_/(1-F_bound_). **Column 5.** Estimated diffusion coefficients for given compounds in free water. ^1^(Fig.5); (Kramer et al., 2007); ^3^(Gendron et al., 2008), describes diffusion coefficients for multiple rhodamine derivatives; ^4^(Gribbon and Hardingham, 1998); ^5^(Waggoner, 1995). **Column 5.** Estimated fold-reduction of diffusion coefficients of unbound compounds in murine myocyte cytoplasm versus water.

As shown in column 2, the fluorescent PEG probes show the least binding to cytoplasmic constituents with bound fractions ranging from 0.12 to 0.52. As expected, the bound fraction of sulforhodamine is high, amounting to 0.95, and the bound fractions of the other fluorophores range from 0.31 to 0.55. Column 3 of Table 1 gives our best estimates of the apparent diffusion coefficients of the fluorophores cytoplasm, calculated from the time constants given in Fig. 5 and in Sup.Fig. S1 and average myocyte length (0.012 cm):

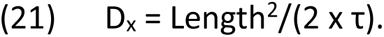

Finally, column 4 gives our best estimates of the diffusion coefficients of free fluorophores after compensating for the bound fluorophore fraction:

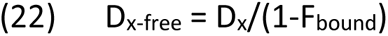

The values range from 0.28 x 10^-7^ cm^2^/s for Bodipy-FL-ATP, very close to the value of 0.4 x 10^-7^ cm^2^/s determined previously (Illaste et al., 2012), to 1.2 x 10^-7^ cm^2^/s for carboxyfluorescein.

### Unrestricted Na diffusion in murine myocytes

Figs. 8 to 11 extend our previous functional analysis of diffusion in murine cardiac myocytes (Lu and Hilgemann, 2017). Fig. 8 illustrates the time course with which Na is lost from patch clamped myocytes after Na is loaded via sarcolemmal NaV channels. Panel A illustrates a protocol in which myocytes were initially patch clamped with NMG as sole monovalent cation in both the cytoplasmic (patch pipette) and the extracellular solutions employed. The extracellular solution contains additionally 7 mM Na to suppress pump current that is generated by contaminating K in standard solutions (Lu et al., 2016). To activate Na/K pump current, the 7mM Na is substituted for 7mM K. Initially, this protocol generates no detectable pump current, indicating that cytoplasmic Na was effectively depleted from the myocytes. Subsequently, the NaV channel opener, veratridine (3 μM) (Zong et al., 1992), was applied together with 120 mM extracellular Na. An inward Na current of about 1 nA developed routinely within 10 seconds and decayed by about 15% over 80 s. When Na and veratridine were subsequently removed, an outward current of ∼90 pA, developed and decayed with a time constant of about 5 seconds. This current likely represents reverse Na current through NaV channels, and the time course likely reflects veratridine dissociation from cardiac myocyte NaV channels.

**Figure 8.**
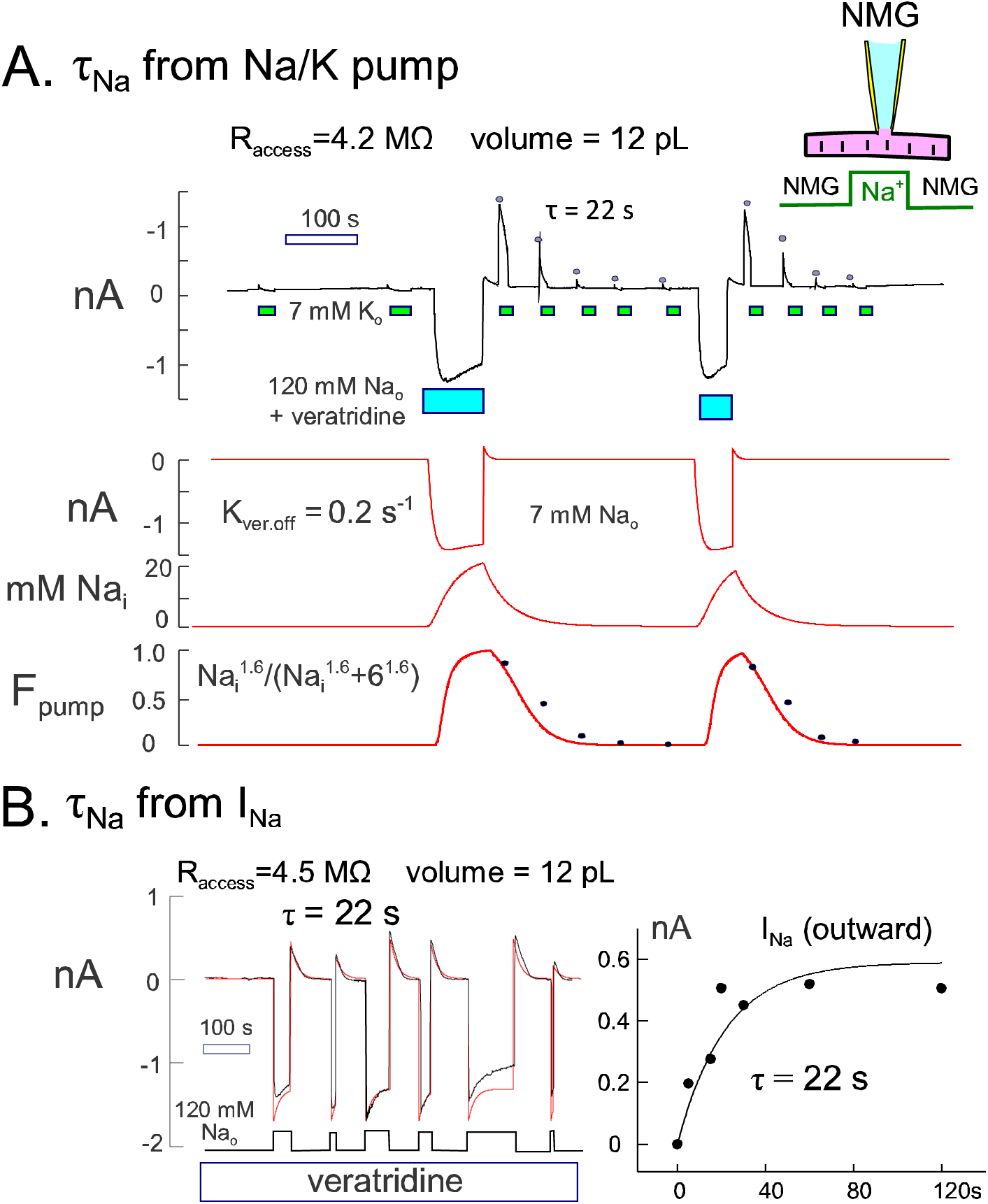
Na exchange in patch clamped murine myocytes using veratridine to induce a large Na influx. **A**. Na exchange estimated from Na/K pump currents. 120 mM NMG-Aspartate on both sides. The extracellular solution contains additionally 7 mM Na or 7 mM K. NMG-Aspartate is exchanged for Na-Aspartate with 3 uM veratridine on the outside. The ∼1 nA inward Na current decays partially, followed by generation of transient outward current upon removal of Na and veratridine. Thereafter, large Na/K pump currents are activated by exchanging 7 mM Na for 7 mM K, and peak currents decay with a time constant of 22 s. Red curves show simulations of Na exchange to a 12 pL cytoplasmic volume. Veratridine dissociates at 0.2 per s, and pump current is proportional to a Hill equation with a slope of 1.6 (Zong et al., 1992). Peak Na pump current magnitudes determined experimentally are plotted with the simulated pump availability. **B**. Na exchange estimated from veratridine-activated Na current. In the presence of veratridine (3 μM), inward Na currents decay partially and outward (reverse) Na currents decay completely with a time constant of 22 s after Na removal. Peak reverse currents after removing Na are plotted in the right panel in dependence on the time that Na was applied. The time constant is also 22 s. Simulations assume that Na turnover is determined solely by the pipette access resistance.

Immediately after removal of Na and veratridine, a large 1 nA Na/K pump current can be activated by applying extracellular K in the same manner as before Na loading. K was applied and removed 4 additional times to determine the time course with which Na is lost from the myocyte via the patch pipette. From these records, we determined the magnitudes of peak pump currents at the 9 points, as indicated, and the corresponding times after removal of Na and veratridine. The peak currents were fit to a single decaying exponential, and the time constant of current decay was determined by least-squares fitting to be 22s. This value is very similar to results for other protocols used to estimate the time course of Na equilibration with patch pipettes (Lu and Hilgemann, 2017). We note that the more rapid decay of Na/K pump current that occurs during application of extracellular K probably reflects an auto-inactivation mechanism of murine cardiac Na/K pumps (Lu and Hilgemann, 2017).

The red curves below the current trace in Fig. 8A are a simple reconstruction of Na homeostasis in these experiments. It is assumed that Na equilibrates instantly within the entire cytoplasm and that Na exchange with the pipette tip occurs with a time constant of 22 s. This time course is accurately predicted when the pipette access resistance of 4.2 MΩ is simulated as a nonselective conductance and the myocyte mixing volume is assumed to be 12 pL (Lu and Hilgemann, 2017). It is assumed that Na escapes from the cell exclusively via the patch pipette tip opening. Although no Na pumping is included, Na/K pump availability is simulated by a Hill equation with a half-maximal Na concentration of 6 mM and a Hill coefficient of 1.6, as defined experimentally for this condition without cytoplasmic K (Lu et al., 2016). Finally, we simulated veratridine binding and dissociation from the myocyte to occur with a time constant of 5 s. With these simple assumptions, the simulation reproduces the experimental results with good accuracy.

Figure 8B illustrates a second protocol that should reveal subsarcolemmal Na accumulation, if it occurs. In this protocol, veratridine was applied continuously in the presence of NMG as sole monovalent cation inside and outside the myocyte. Then, Na (120 mM) was applied instead of NMG on the outside, and inward Na currents of about 1.5 nA were generated. The inward current decays by 15%. The average time constant, determined by fitting the individual current records to single exponential functions was 22 s. The outward current which develops upon removal of extracellular Na also decays with a time constant of 22 s. As shown in the right panel of Fig. 1B for repeated Na applications, removal of Na at different times verifies that the outward Na current develops with the same time course as the inward current decays. The red curves in panel B of Fig. 8 show a simple reconstruction of Na homeostasis in this experiment, assuming that Na equilibrates instantly within the myocyte and exchanges to the patch pipette with a time constant of 22 s from a mixing volume of 12 pL. The results give no evidence for complexities beyond those represented.

In Sup. Fig. S2 we describe similar experiments using myocytes loaded with CoronaGreen AM Na-selective dye (5 μM for 30 min at 25°C) (Iamshanova et al., 2016). CoronaGreen fluorescence changes in similar experiments (n>20) were in general of small magnitude in relation to signal drift and bleaching We provide an example in which fluorescence changes seemed reliable. They occurred with a time constant of 28 s.

### Restricted ATP diffusion in myocytes

Figures 9 to 11 provide evidence that diffusion of ATP and ATP analogues in murine myocytes is substantially restricted in comparison to Na. Fig. 9A presents functional results for an ATP analogue, AMPPNP, which inhibits the Na/K pump by competing with ATP (Beaugé and Glynn, 1980; Robinson, 1980). To determine the likely time course of diffusion of AMPPNP from the pipette tip to the sarcolemma, we employed a fast pipette perfusion technique to inhibit and reactivate Na/K pump current by infusion and removal of AMPPNP (Hilgemann and Lu, 1998). The experimental record is a solid black line, and single exponential functions fitted to current changes are given in gray. At the onset of the experiment, pump current was activated and deactivated by application and removal of extracellular K (7mM) in the presence of 80 mM Na and 4 mM MgATP in the patch pipette. Subsequently, the pump current was activated and left activated with 7 mM extracellular K. After 30 s, pipette perfusion of 2 mM AMPPNP in the continued presence of 4 mM ATP inhibited the pump with a time constant of 25 s. Thereafter, AMPPNP-free solution was perfused into the pipette tip, and pump current was restored with a time constant of 160 s. As illustrated further, results could be repeated with good reproducibility.

**Figure 9.**
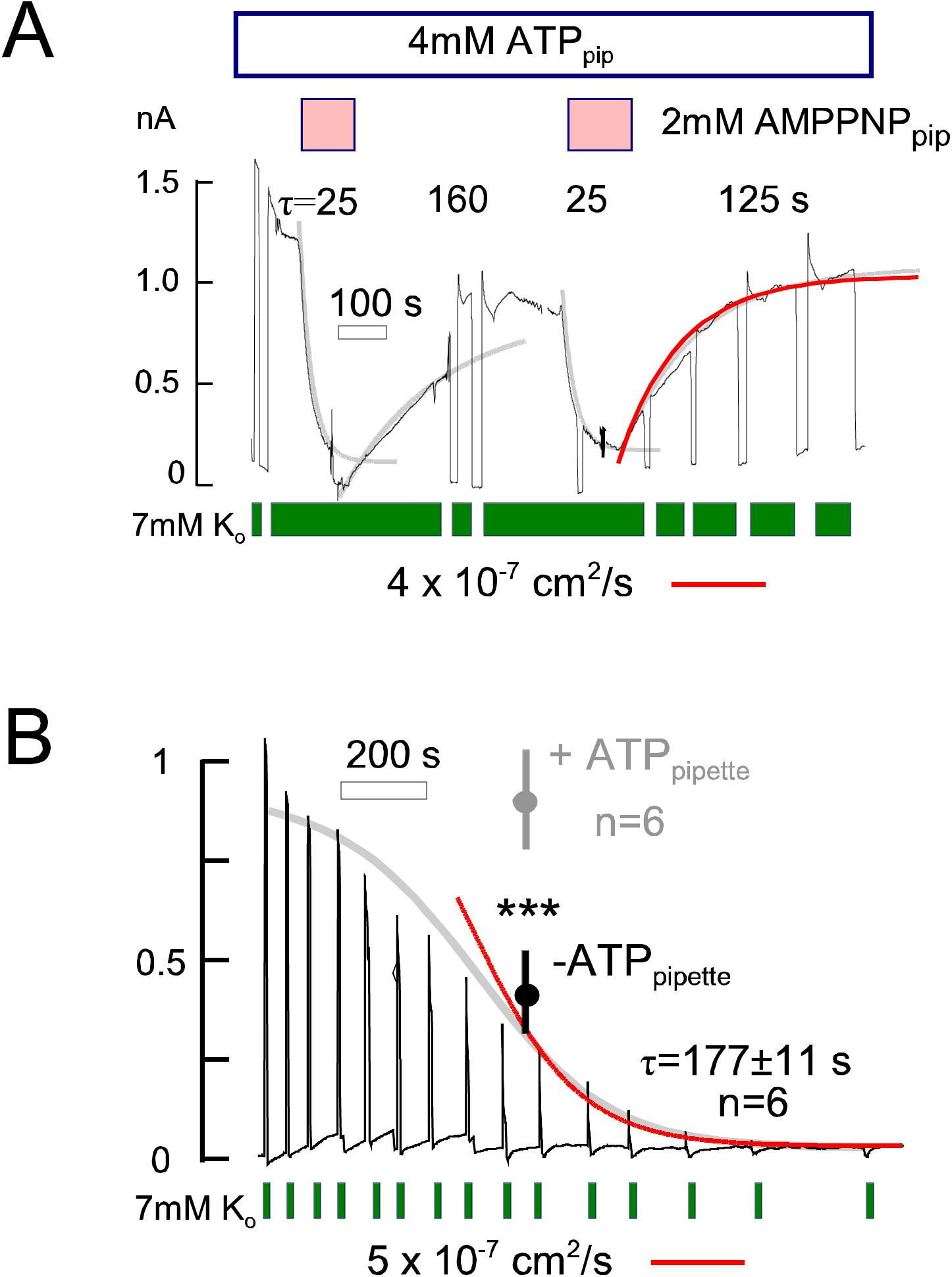
Functional analysis of nucleotide exchange between murine myocyte cytoplasm and patch clamp pipette tip. **A.** Pipette perfusion of AMPPNP. A rapid pipette perfusion technique (Hilgemann and Lu, 1998) was used to apply and remove nonhydrolyzable ATP, AMPPNP, from the cytoplasmic side. The pipette contained 4 mM ATP and 80 mM Na throughout. During continuous pump activation with 7 mM K, AMPPNP (2 mM) perfusion causes pump shut down with a time constant of 25 s. Pumps reactivate with a time constant of 160 s (gray curves) during AMPPNP removal, the protocol repeats accurately. Gray curves are single exponential functions, fit to results. The red curve in panel A simulates a solute with a diffusion coefficient of 4 x 10^-7^ cm^2^/s through a 120 μM long cylinder, open at one end. **B.** Pump current changes in the presence and in the absence of ATP in the patch pipette. Pump currents were activated for just 2 s to ensure that decay reflects exchange of Na to the pipette, rather than Na depletion via pump activity or Na/K pump inactivation. Mean values with standard errors are given for current magnitudes at 10 min. With ATP, currents decreased by only 10%. Without ATP, Students T-test verified that a 65% decrease of pump current at 10 min was highly significant with respect to ATP containing solution. In all experiments without ATP, pump current decayed completely over 15 min. The final phase of current decay had a time constant of 177 s on average (n=6), and the entire time course of decay is simulated (gray curve) assuming that ATP activates Na/K pumps with a K_50_ of 80 μm and diffuses longitudinally with a diffusion coefficient of 5 x 10^-7^ cm^2^/s.

The fact that pump current is strongly inhibited by 2 mM AMPPNP suggests that the affinity of AMPPNP for the murine Na/K pump is quite high. Alternatively, the cytosolic ATP concentration might be less than the concentration employed in the pipette (e.g. as a result of fast ATP consumption versus synthesis). In both cases, pump inhibition will occur more rapidly than AMPPNP concentration changes at the sarcolemma. The time course with which pump current is restored on removing AMPPNP should however represent accurately the time course with which AMPPNP out of the myocyte. Pump reactivation begins with very little delay, and the red curve in Fig. 9A, given with the final recovery response, accurately reconstructs the results. This simulation assumes that pump activity is proportional to the average concentration of a solute in a cylinder 120 micron long. Starting with a uniform concentration, the solute diffuses out of the cylinder at one end with a diffusion coefficient of 4 x 10^-7^ cm^2^/s. zIn other words, the time course of AMPPNP (and ATP) diffusion out of myocytes is likely dictated mostly by diffusion through the cytoplasm, rather than by access resistance of pipette tips.

Figure 9B shows the time course with which pump current declines when ATP-free pipette, solution is employed from the start of experiments with 80 mM Na in the pipette solution (n=6). As indicated by the error bar at the 10 min time point, pump current is stable when 4 mM ATP is included in this pipette solution. Without ATP, pump current declines in a sigmoidal fashion over time, and the final phase of current decay is approximately exponential with an average time constant of 177 s (n=6). The gray line in Fig. 9B is a simulation assuming that ATP diffuses out of a 120 micron long cylinder, open at one end, with a coefficient of 5 x 10^-7^ cm^2^/s, starting from a uniform concentration of 4 mM. and activating Na/K pumps with a half-maximal concentration of 80 μM, as determined excised cardiac giant patches (Collins et al., 1992).

Figures 10 and 11 describe experiments to assess the diffusion of anions into murine myocytes via their coupling to Na diffusion, as outlined in Methods (Fig. 3). Fig. 10 presents simulations of the experiments in which we attempted to follow the diffusion of 120 mM Na(Mg)ATP into myocytes through the pipette tip, followed by depletion of Na and MgATP during the activaiotn of Na channels. As described subsequently, we accessed changes of the cytoplasmic Na concentration by applying and removing the Na channel opener, veratridine, in the absence of extracellular Na. For Fig. 10, diffusion was simulated to occur over 110 µm, the pipette opening was assumed to have a resistance 10-times greater than each cross-section of the myocyte, simulated to be 5 µm thick, and the myocyte cytoplasm was assumed to contain 50 mM of fixed negative charges. In Fig. 10A, cytoplasmic Cl and K are 120 and 70 mM at the onset, and the pipette contains 120 mM Na(Mg)ATP with no Cl or K. Thus, on cell opening K and Cl diffuse of of the myocyte with a time constant of about 12 s. At first, Na diffuses into the myocyte rapidly in exchange for K, but after K is lost Na and MgATP equilibrate with the myocyte cytoplasm with the time course of their coupled diffusion constant. When experiments are performed in the absence of extracellular Na, activation of plasma membrane Na conductance will then cause Na depletion that reflects the couple K(Mg)ATP diffusion constant. As shown in Fig. 10 B, however, the initial time course of cytoplasmic Na depletion upon opening Na channels is strongly affected by the presence 5 mM KCl, which we include in the pipette solutions to maintain stable patch clamp via Ag/Cl electrodes. When 5 mM pipette KCl was included in the simulation, Na initially depletes rapidly with the time course of K diffusion, when Na channel are activated.

**Figure 10.**
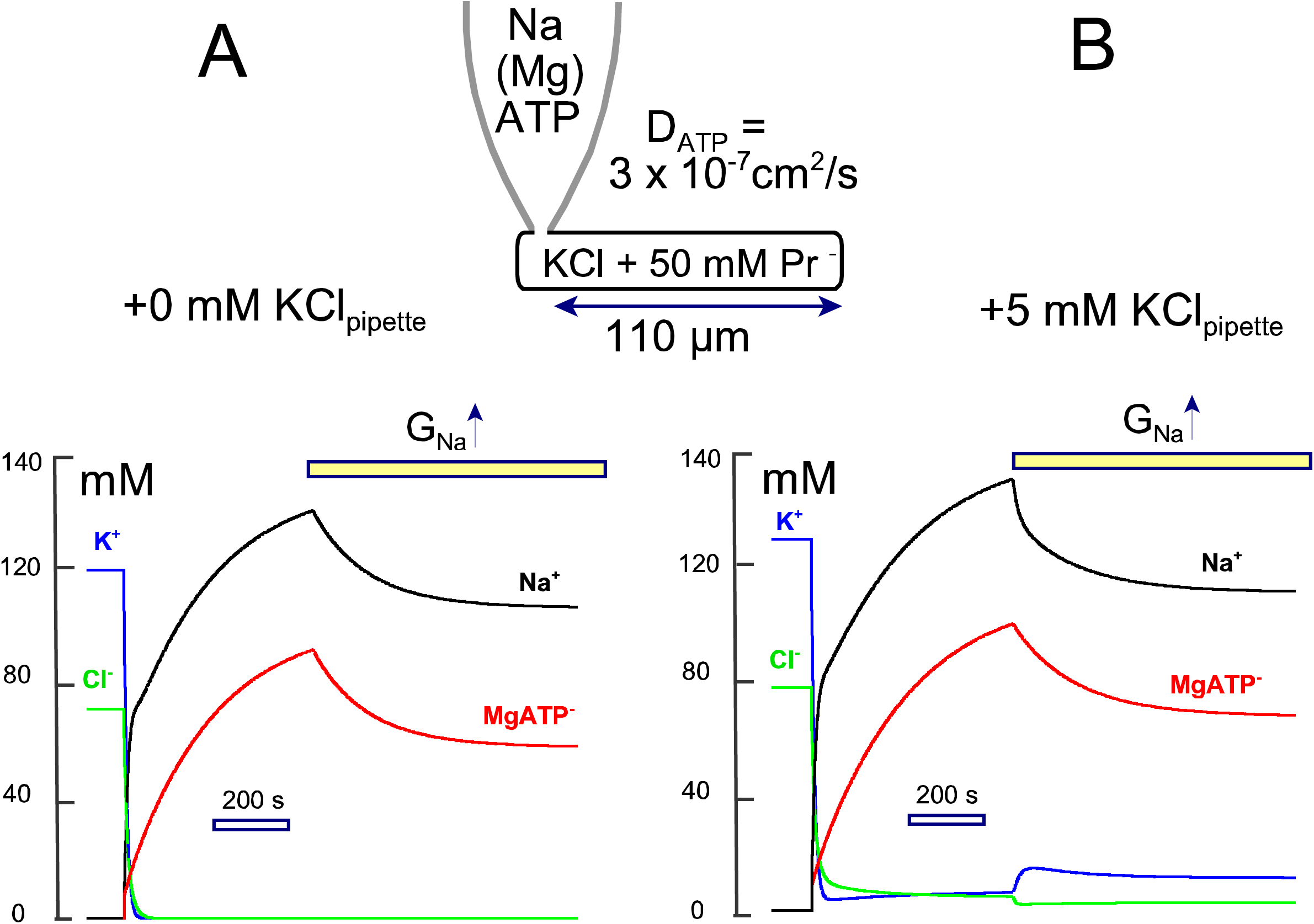
Simulated diffusion between cardiac myocytes and a patch pipette tip with fixed ion/solute concentration. Relative anion diffusion coefficents can be determined from the time course of coupled cation diffusion into cardiac myoyctes. Access resistance waas simulated to be 2-fold greater than the longitudinal myocyte resistance when equilibrated with KCl. **A.** The myocyte initially contains KCl, namely 120 mM K and 70 mM Cl with fixed anionic charges making up the rest of cytoplasmic anions. D_K_ and D_Cl_ were 1.2 x 10^-5^ cm^2^/s. Initially, the pipette contains Na(Mg)ATP (120 mM). D_Na_ is 0.7 x 10^-5^ cm^2^/s and D_ATP_ is 3 x 10^-7^ cm^2^/s. The extracellular solution contains no Na. A surface membrane Na conductance can be activated and deactivated arbitrarily and was set to generate an outward Na current of ∼2.5 nA. Upon opening, KCl is lost rapidly from the myocyte. Thereafter, Na(Mg)ATP diffuses slowly into the myocyte in a coupled fashion, limited by the diffusion coefficient of MgATP. When MgATP is the sole anion present in the cytoplasm, activation of the Na conductance results in coupled depletion of Na and MgATP with essentially identical time courses. **B.** Equivlent simulation in which the pipette solution contains additionally 5 mM KCl. After the initial rapid loss of KCl and gain of Na in exchange for K, Na accumulates in the cytoplasm with close coupling to the accumulation of MgATP. However, Na depletion during activation of Na channels is significantly determined by a rise of cytoplasmic K and loss of residual Cl, in addition to the slower progression of MgATP depletion. Thus, the equilibrtion time courses allow more accurate estimates of diffusion coefficients in these experiments than the depletion time courses.

Figure 11 shows experimental results to evaluate the diffusion of four anions into myocytes via their coupling to Na. Myocytes were initially incubated in a solution containing 140 mM NMG-MES (with 0.5 mM EGTA and 4 mM MgCl2 at pH 7.0) for 15 min. Patch pipettes contained 130 mM Na with 130 mM of a chosen anion, 5 mM KCl, and 1 mM MgATP. Seals were formed without opening the sarcolemma. Then, the myocytes were placed in a solution stream with 120 mM NMG-MES as extracellular solution at 36°C, and whole-cell patch clamp was established via suction. Thereafter, myocytes were intermittently switched for ∼20 s to an extracellular solution containing 25 µM veratridine to open Na channels and activate large outward Na currents. Repeating the application of veratridine at ∼ 1 min intervals, Fig. 10 A illustrates that Na currents were nearly maximal within 40 s of opening myocytes when 120 mM NaCl was employed in the pipette. The outward Na current decayed by about 12% with a time constant of 24 s (see fitted exponential in red). In contrast, Fig. 10 B shows that outward Na current increased over several veratridine applications when 120 mM Na-aspartate was used in the pipette solution. As expected for rather slow diffusion of aspartate, decay of current during veratridine application was larger and proceeded with a longer time constant than with Cl. Using 120 mM Na(Mg)ATP and 130 mM Na with 500 MW carboxy-PEG, these changes became still more pronounced, as illustrated in Figs. 10C and 10D, respectively. Figs. 10D and 10E present the composite results for at least 6 experiments in each condition. The average equilibration and current decay time constants for NaCl were 30 and 15 s, respectively, those for Na-aspartate were 53 and 24 s respectively, those for Na(Mg)ATP were 121 and 40 s, respectively, and those for Na-PEG-COOH were 148 and 84 s, respectively.

**Figure 11.**
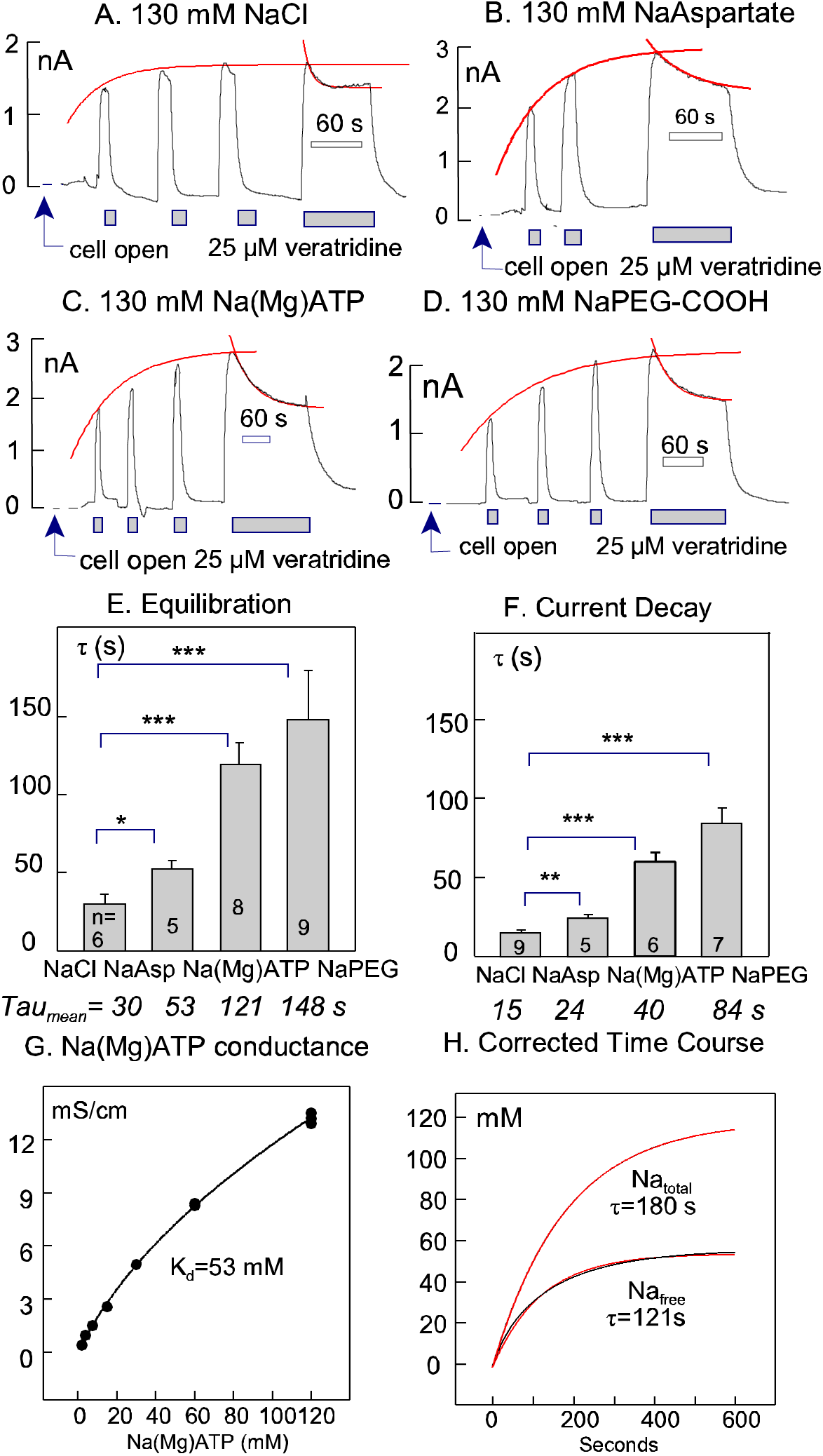
Relative anion diffusion coefficeints determined from time course of couipled Na and MgATP diffusion into murine myocytes. **A-D.** Representative patch clamp records of myocytes opened with pipette solutions containing 130 mM NaCl, Na-aspartate, Na(Mg)ATP, or Na-PEG-COOH (2kD). The pipete also contained 5 mM KCl and 8 mM HEPES titrated to pH 7.0 with KOH. The extracellular solution contained 130 mM NMG-aspartate, 4 mM MgCl_2_, and 10 mM HEPES titrated to pH 7.0 with NMG. As indicated, 10 µM vertridine was applied and removed for a few seconds, long enough to achieve a peak outward Na current. The final veratridine application was continued for long enough to induce current decay, indicative of cytoplasmic Na depletion. The time course with which peak veratridine currents approached a maximun was fitted to a single exponential function. **E.** Composite results for the rising peak veratridine currents after myocyte opening. The mean time constants for 5 to 9 measurements are given below the respective columns for the four ion pairs examined. **F.** Composite results for the decay of currents during continued veratridine application. **G.** Concentration-conductance relation of Na(Mg)ATP solutions, determined as in Figs. 2A-C. The best least-squares fit to equation #1 has a dissociation coefficient of 53 mM. H. Corrected time course of Na accumulation during patch clamp with 120 mM Na(Mg)ATP. The time constant with which Na currents saturate was assumed to be 121 s, as determined experimentally (lower curve). From this time course, reflecting free Na changes, the time course with which total Na equilibrates is calculated from equation #1 after rearranging to calculate the total ion pair concentration, ‘C’ (upper curve, together with least squares fit to an asymptotic exponential function).

To evaluate results for Na(Mg)ATP, it must additionally be taken into account that Na(Mg)ATP at a concentration of 130 mM does not fully dissociate, as already described for K(Mg)ATP in Fig. 2C. Fig. 11G shows the relevant concentration-conductance relation for N(Mg)ATP, which is best described by a dissociation constant of 53 mM. As illustrated in Fig. 11H, this relationship affects the time constant of Na current equilibration, causing an underestimation of the time constant for Na equilibration. When results are corrected for the tendency of free Na to saturate with increasing concentration, the time constant for total Na equilibration is 180 s, closely consistent with results of Fig. 9 for ATP and AMPPNP diffusion. In conclusion, these experiments lend support to the estimation that MgATP diffuses with a coefficient of about 4 x 10^-7^ cm2/s in murine cardiac myocytes. Importantly, the equilibration time constants increase with increasing molecular weight of a larger extent than occurs in free water. In other words, the diffusion barriers in myocyte cytoplasm have a pore-like barrier function.

### Diffusion restriction by internal membranes versus myofilaments

A key question posed by these results is whether diffusion restrictions arise primarily from networks of myofilaments or membranes in murine myocytes. To address this question, we analyzed diffusion through permeabilized and physically confined murine cardiac myocytes, as introduced in Methods and described in detail in Fig. 12. Briefly, thick-walled, bullet-shaped glass pipette tips were formed, as used for the giant patch technique (Hilgemann and Lu, 1998), with opening diameters of 8 to 12 microns. With these dimensions, myocytes were readily aspirated into the tips, and intact myocytes formed seals with resistances ranging from several 10’s of megohm to gigaohm values (see Fig. 11 A). A background solution containing 20 mM KCl was employed, and in this configuration the application of 50 mM of the additional ion pair, NaCl or K(Mg)ATP, increased the tip conductance by a factor of 3 to 5 (see leftward half of the conductance trace). Subsequently, application of 90 µM of β-escin to permeabilize the sarcolemma (Konishi and Watanabe, 1995) induces an approximately 8-fold increase of the tip conductance. Thereafter, application of 50 mM NaCl or K(Mg)ATP cause rapidly reversible 3-fold increases of conductance. While the peak conductances for NaCl and K(Mg)ATP were similar, the time courses of conductance changes upon changing solutions were distinctly slower for K(Mg)ATP in comparison to NaCl. This is quantified from the first derivative of the conductance record, shown below in Fig. 12A. We note that the time courses of conductance changes before permeabilization are too fast to quantify accurately, although peak velocities were higher for NaCl than for K(Mg)ATP. After permeabilization, the time constants of conductance changes can be accurately fit to single exponentials. The average time constants of rising and falling NaCl conductances are 0.27 versus 1.7 s for K(Mg)ATP, and the ratio of peak velocities for the two ion pairs is 2.3:1. As illustrated below the experimental records, the ion coupling principle elaborated in Fig. 2 indicates that the MgATP diffusion coefficient is about 10% of the Cl diffusion coefficient, in good agreement with results of Figs.

**Figure 12.**
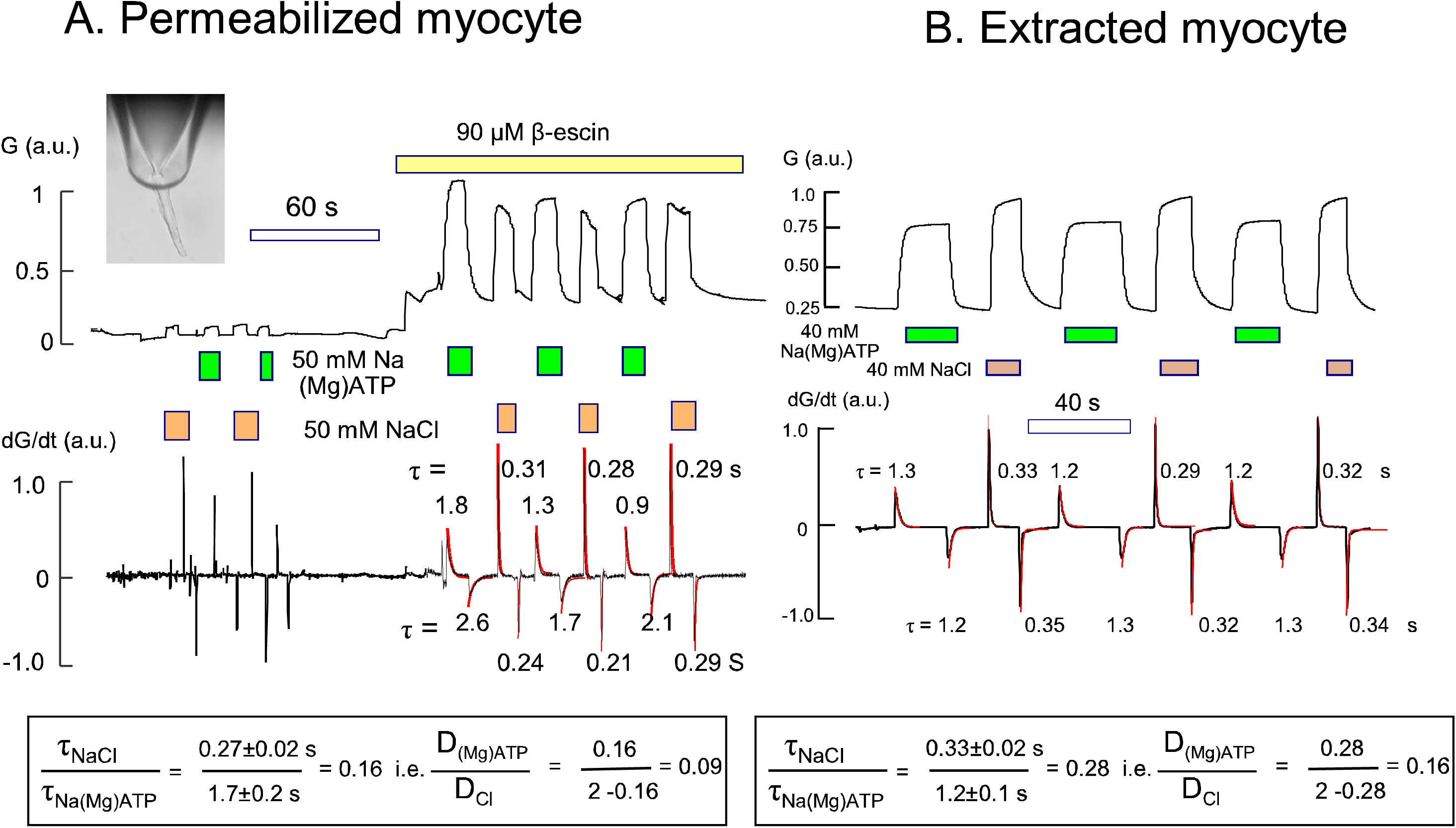
Coupled ion diffusion in PM-permeabilized and extracted murine cardiac myocytes. A. Myocytes were aspirated into the tip of a highly polished pipettes, as shown in the inset, employing a background solution that cotained 20 mM KCl. The sarcolemma was then permabilized by application of 80 µM β-escin, and thereafter conductances induced by applying 50 mM NaCl verus 50 mM Na(Mg)ATP are compared. From the first derivative of the conductance signal, the maximal rates and time constants of conductance changes were determined. Time constants for NaCl signals were 16% of those for Na(Mg)ATP. From equation #8, this indicates that the diffusion coefficient of MgATP is 9% of the Cl diffusion coefficient. B. Equivalent experiments to Panel A were performed with myocytes that had been extensively membrane-extracted, as described, with glycertol and Triton X-100. The ratios of diffusion coeffients for MgATP and Cl were nearly twice as large, indicating that MgATP diffusion is less restricted in extracted myofilaments than in permeabilized myocytes.

To determine the role of membranes in the results of Fig. 11A, membranes were aggressively extracted by incubation for 1 to 7 days at 4°C in an extraction solution containing 50% glycerol, 25 mM KCl, 10 mM EGTA, 0.5 mM MgCl2 and 2 mM Triton X-100 (Palmer et al., 1996). At low-magnification (20x), these myocytes were indistinguishable from the permeabilized myocytes, as in Fig. 13A. Nevertheless, as shown in Fig. 12B, the velocities with which conductances changed in equivalent solution switch experiments were significantly faster. Most importantly, the ratio of the apparent MgATP diffusion coefficient to Cl diffusion coefficient was 0.14 in extracted myocytes compared to 0.09 in permeabilized myocytes. Accordingly, intracellular membranes evidently account for at least one-half of the diffusion restriction encountered by solutes in murine myocytes.

**Figure. 13.**
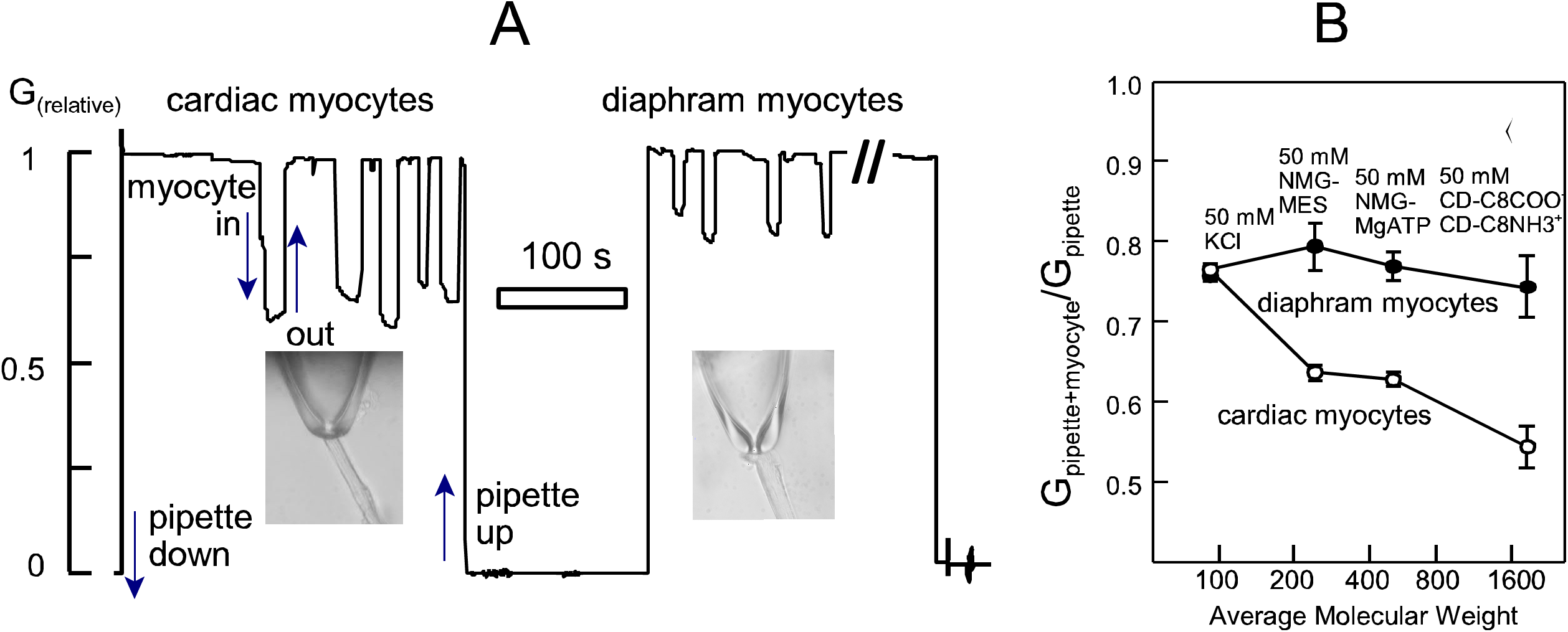
Conductance restrictions induced by cardiac and diaphragm myofiliments and their dependence on solute molecular weight. **A**. Representative experiment with membrane extracted myoctyes using a solution of 50 mM NMG-MES, 0.5 mM MgCl_2_, 0.5 mM EGTA, and buffered to pH 7 with 4 mM HEPES. Initially, 5 cardiac myocytes were aspirated and released from the pipette tip, cardiac myocytes were replaed with diaphram myocytes in the dish, and then 4 diaphram myocytes were aspirated and released. As apparent in the conductance record, conductances with aspirated cardiac myofilaments were substantilly less than with aspirated diaphram myofilaments. **B.** Comparison of conductance changes caused by cardaic and diaphram myofilaments for 4 ion pairs. For all results, n>6. The graph plots the ratio of pipette conductance in the presence versus absence of aspirated myofilaments. Conductances of cardiac and diphram myofilaments were similar with 50 mM KCl. While cardiac mhyofilaments induced progressively larger decreases of conductance with ion pairs of increasing molecular weight, diaphram myfilaments caused similar conductance reductions for all four ion pairs employed, namely KCl, NMG-MES, NMG-(Mg)ATP, and β-methylcyclodextrin complexes with octanoate and octylamine.

### Cardiac myofilaments restrict diffusion more than skeletal myofilaments

To better understand the role of myofilaments in diffusion restriction, we determined for 8 ion pairs to what extent their immediate conductivity was decreased by the presence in the pipette tip of either extracted cardiac myofilaments or extracted diaphragm (skeletal) myofilaments. Fig.13A shows a typical experiment in which conductance measurements were carried out continuously with a single pipette using 50 mM NMG-MES as the background solution. Five cardiac myocytes were sequentially aspirated and ejected from the pipette tip. Then, murine diaphragm myocytes were added to the chamber and 4 extracted diaphragm myocytes were aspirated and ejected. The average drop of conductance upon aspirating cardiac myocytes is significantly greater (p<0.01) than the average change by skeletal myocytes. Fig. 13B shows results for 8 ion pairs, plotting the average molecular weight of the anion and cation on the X-axis and the pipette conductance with aspirated myocyte as a fraction of the conductance without myocytes. From left to right, the results show that for 50 mM KCl both cardiac and diaphragm myocytes decrease pipette conductance by 24%. This is consistent with hydrated myofilaments making up about 20% of the myocyte volume (Lu and Hilgemann, 2017). We mention that for 140 mM KCl solution, the drop of conductance was increased 10% for cardiac filaments and 5% for diaphragm filaments, as expected if ions promote myofilament interactions (not shown). Using ion pairs of increasing molecular weights at a concentration of 50 mM, the decrease of conductance induced by cardiac myofilaments shifts increases from 24% to 45%, while the restriction imposed by skeletal filaments is nearly constant at 25%. The largest ion pair was made using 50 mM β-methylcyclodextrin complexed with octanoate and octylamine, respectively. Stable monovalent complexes were verified by the fact that osmolarity of the final solution with 50 mM of both complexes was 104 mosmol/L.

Figure 14 compares results for further conductance experiments with cardiac myofilaments with results for intact cardiac myocytes. Results in Fig. 14 are plotted in dependence on calculated hydrated radii (https://www.fluidic.com/toolkit/hydrodynamic-radius-converter/). For conductance measurements the average hydrated radius was employed, and results were then fitted to Boltzmann functions of the form,

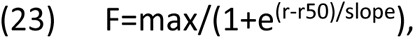

**Figure 14.**
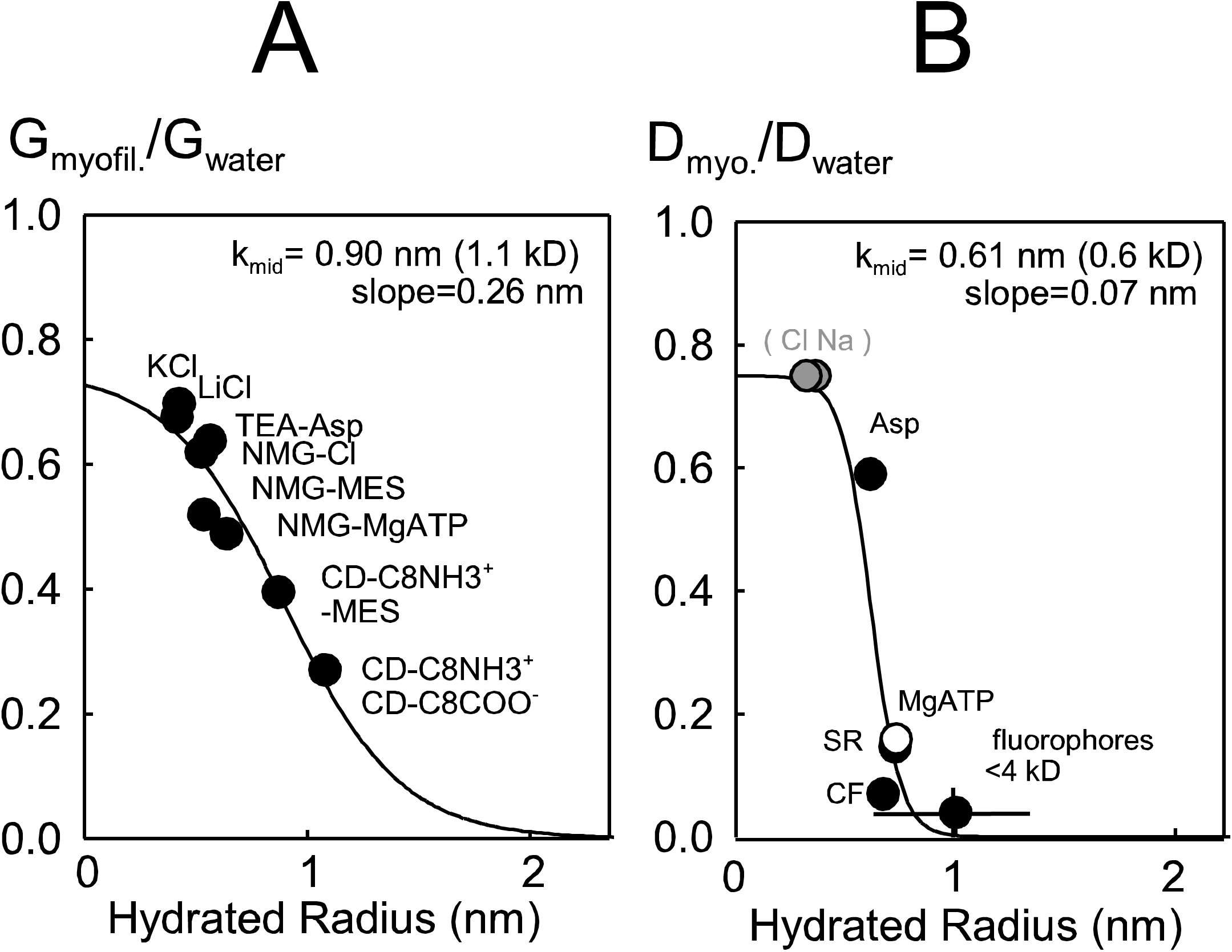
Comparison of diffusion restrictions employing murine cardiac myofilaments versus intact murine cardiac myocytes, plotted in dependence on average hydrated radius of the ion pairs **A.** Ratio of pipette tip conductance with and without aspirated cardiac myofilaments for 8 ion pairs. **B.** Ratio of best estimates of diffusion coefficients for selected solutes in murine BL6 myocytes versus free water. Results are fitted to Boltzmann equations, the slope of which is three-fold steeper for intact myocytes.

where F is the fractional value with respect to free water. For the extracted myocytes employed in Fig. 14A, eight different ion pairs were employed with at least 10 measurements per pair, and the standard errors of the results were within the diameter of the data points plotted. The water/myofilament conductance ratio falls off with mid-point of 0.26 nm, corresponding to a molecular weight of 1.1 kD, and a slope of 0.26 nm. Fig. 14B summarizes our major results for intact myocytes, plotting the ratio of the estimated diffusion coefficients in myocytes to that in water. We assumed that a maximal ratio of 0.75, corresponding to the percent of cytoplasmic space taken up by myofilaments. Cl and Na are assigned values of 0.75 because we obtained no evidence that their diffusion is restricted. Results of Fig. 11 indicate that aspartate diffuses with a coefficient one-half that of Cl. Thus, aspartate is only marginally restricted in myocytes. Figs. 9 to 12 indicate that MgATP diffuses between 8- and 10-fold slower in cytoplasm than in free water, whereas fluorophores with molecular weights of 450 to 4000 all diffuse more than 10-times slower than in free water, some as much as 50-times slower. Thus, the Boltzmann function has a mid-point of 0.61 nm (∼0.6 kD) and a steep slope factor of 0.07 nm. Equivalent results for 10% gelatin are provided in Sup. Figs.5-10, including ATP diffusion measurements via luciferase luminescence, analysis of ATP and ADP diffusion via conductance measurements, and description of the break-down of diffusion restriction that occurs when gelatins are depolymerized at 37°C.

### Evidence for a role of mitochondria in restricting diffusion in the murine myocyte cytoplasm

Finally, we designed experiment to test whether internal membranes, specifically mitochondrial membranes, play a significant role in restricting diffusion in murine cardiac myocytes. We reasoned that the outer mitochondrial porin channels would influence diffusion if mitochondrial networks in myocytes indeed form significant diffusion barriers. Specifically, we hypothesized that porins would facilitate long range diffusion by facilitating solute passage through intermembrane space of mitochondria. Accordingly, we analyzed myocytes from VDAC1 knockout mice. As noted in Methods, breeding of VDAC1 knockout animals requires a mixed genetic background, and we employed a CD1/J6/129svJ background. We then examined both the decline of Na/K pump currents during patch clamp with ATP-free pipette solution, and we examined the diffusion of carboxyfluorescein into patch-clamped myocytes.

Fig. 15A shows examples of pump current run-down in WT CD1/J6/129svJ cardiac myocytes and in VDA1 knockout myocytes from CD1/J6/129svJ mice. Surprisingly, as shown in Fig. 15B, the decay of pump currents was more than 2-fold faster in myocytes from CD1/J6/129svJ mice than in myocytes from WT BL6 mice, and similar results were obtained from mice of both sexes. In addition, pump current run-down was slowed by more than two-fold in VDAC2 knockout myocytes compared to control CD1/J6/129svJ myocytes. Accordingly, pump run-down was similar in CD1/J6/129svJ VDAC1 knockout mice and in myocytes from WT BL6 mice. We further report that pump run-down in CD1/J6/129svJ myocytes was similar with 80 or 20 mM cytoplasmic (pipette) Na.

**Figure 15.**
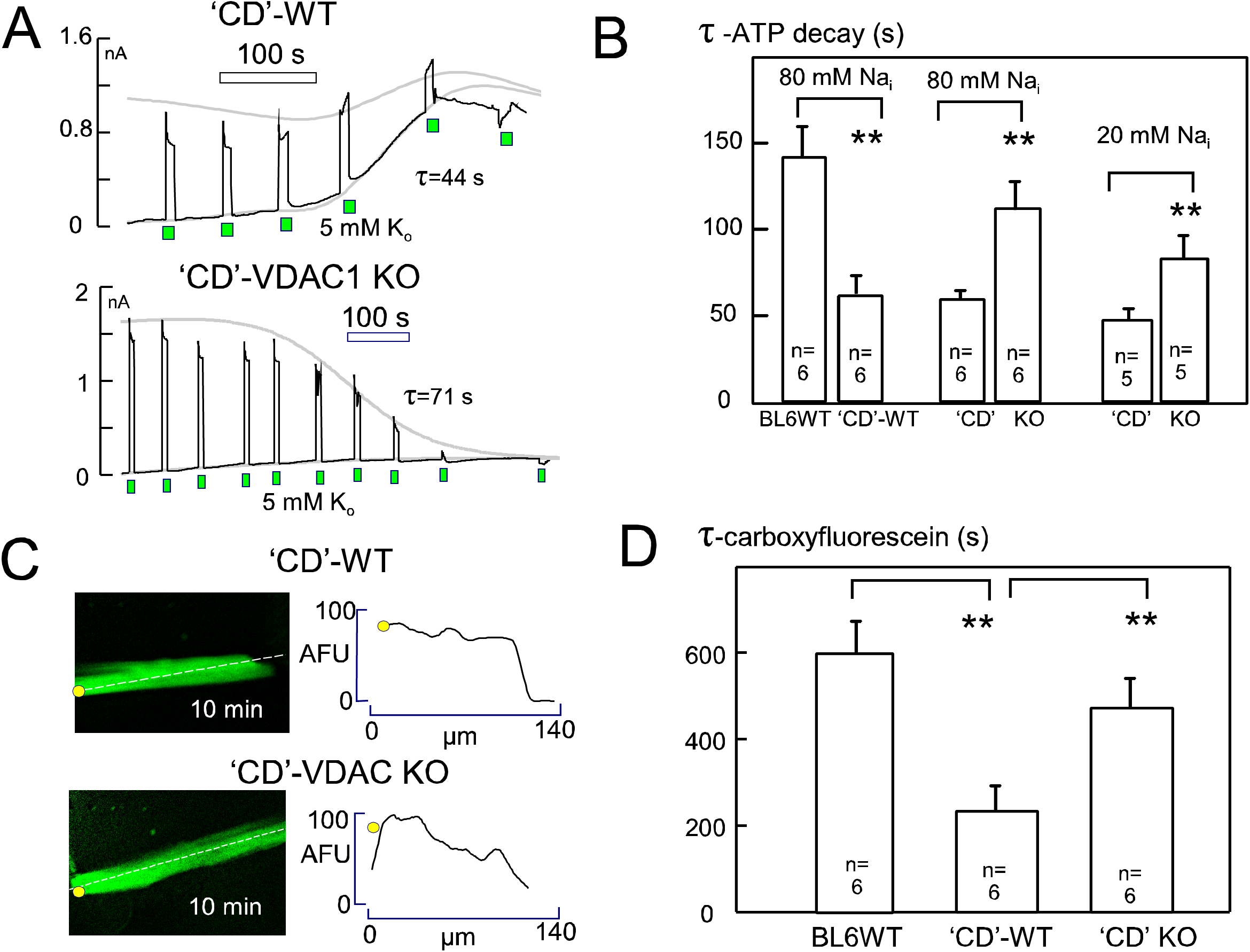
Diffusion restriction in murine cardiac myocytes depends on strain and is enhanced by deletion of VDAC1 porin channels. ‘CD’ indicates myocytes from CD1/J6/129svJ mice. **A.** Representative records of Na/K pump current decline using patch pipettes without MgATP for ‘CD’ wild-type and ‘CD’/VDAC1-defidient myocytes. **B.** Composite results. Decline of Na/K pump current is more than two-fold faster in ‘CD’ wild-type myocytes than in BL6 wild-type myoyctes. Pump current decline is two fold slower in ‘CD’ myocytes lacking VDAC1 than in ‘CD’ wild-type myocytes. Results were similar using either 80 or 20 mM cytoplasmic Na. **C**. Micrographs of ‘CD’ wild-type and VDAC1-deficient myocytes after 10 min of patch clamp with 10 µM carboxyfluorescein. The longitudinal fluorescence gradient is still pronounced in the myocyte lacking VDAC1 after 10 min. **D.** Similar to results for Na/K pump currents, the time constant with which carboxyfluorescein equilibrates in BL6 wild-type myocytes is more than two-fold greater than in ‘CD’ wild-type myocytes, and the time constant is more than two-fold greater in’CD’ myocytes lacking VDAC1 than in wild-type ‘CD’ myocytes.

As shown in Figs. 15C and D, results for diffusion of carboxyfluorescein into myocytes were similar to results for Na/K pump run-down. Diffusion of carboxyfluorescein into myocytes from CD1/J6/129svJ mice was 2.5 fold faster than for myocytes from BL6WT mice, and diffusion of carboxyfluoescein into VDAC1 knockout myocytes was >two-fold slower than for myocytes from WT CD1/J6/129svJ myocytes. One simple explanation for slower diffusion rates in BL6 versus CD1/J6/129svJ myocytes would be the expression level of VDAC channels. Therefore, we tested by Western blotting whether expression levels of VDAC isoforms might be different in the different mice strains. As shown in Sup. Fig. S3, VDAC1 was entirely absent in the knockout myocytes, and expression levels of other VDAC channels was not significantly different in BL6 versus WT CD1/J6/129svJ myocytes.

## Discussion

In this article we have used patch clamp and fluorescence microscopy to characterize the diffusion of Na, ATP, diverse fluorophores, and a series of PEGs in cardiac myocytes. We have compared, as possible, results from intact myocytes to results employing PM-permeabilized myocytes and membrane-extracted myocytes (i.e. myofilaments). In Supplemental Data we also provide comparable measurements of diffusion in gelatins, prepared at concentrations relevant to the protein content of cardiac myocytes, and we describe experiments with macromolecules that do not form gels.

Diffusion coefficients fall off steeply in intact cardiac myocytes over a molecular weight range of 300 to 1000 (Fig. 14), and diffusion coefficients of many commonly employed fluorophores are more than 30-fold reduced in the BL6 cardiac myocyte cytoplasm compared to water. Evidently, both cardiac myofilaments and mitochondria contribute to diffusion restrictions. Extracted cardiac myofilaments clearly restrict diffusion considerably (Fig.12). Potentially, our measurements underestimate the restrictions, since myofilaments tend to swell subsequent to membrane extraction. However, it is also certain that deletion of the outer mitochondrial porin, VDAC1, significantly reduces solute diffusivity in murine myocytes (Fig 15). Clearly, therefore, diffusion restrictions in murine myocytes arise from both myofilaments and mitochondrial networking (Piquereau et al., 2013). The contribution of outer mitochondrial porin channels to these restrictions suggests that porins themselves promote the steeper decline of diffusivity with increasing molecular weight in intact versus extracted myocytes (Fig. 14).

Although porins contribute to diffusion restrictions in myocytes, we document in Supplemental Data that gelatins generate diffusion restrictions with characteristics similar to those of intact BL6 myocytes. Restrictions in BL6 cardiac myocytes are greater than described for both skeletal muscle (Kushmerick and Podolsky, 1969) and myotubes (Arrio-Dupont et al., 1996). It is impressive in this light that cells with one of the highest metabolic rates of any cells in biology, murine cardiac myocytes (Naumova et al., 2003), also appear to have cytoplasmic architectures that are unusually restrictive to solute diffusion. We hypothesize that, as a result of combined myofilament and mitochondrial diffusion barriers, solutes larger than monovalent ions will tend to diffuse through intermembrane mitochondrial spaces where diffusion may be less restricted than in the myofilament space.

### Na diffusion in cardiac myocytes

The idea that local Na gradients modify Na transport function gained much support over several decades (Lu and Hilgemann, 2017). In one recent study cytoplasmic Na accumulation was promoted in mouse cardiac myocytes by rapid depolarizations that activated voltage-gated Na channels (Skogestad et al., 2019). Na/K pump currents were reported to be enhanced by an amount that could not be explained if Na influx was occurring into the entire cytoplasm, and it was concluded that Na accumulates in a subsarcolemmal compartment with turnover to bulk cytoplasm occurring over 1 to 2 minutes, the time course over which Na/K pump currents decayed.

Accordingly, we devised experiments, based on the same principles, to examine the influence of Na influx via NaV channels on subsequent Na/K pump activity (Fig. 8A). To do so, we employed Ca-free and essentially Na-free conditions, and we activated large, well-defined Na currents by application of extracellular Na and veratridine together. These currents could be rigorously quantified to know how much Na entered the myocytes. Subsequent Na/K pump currents decayed back to negligible values with a time constant of 22 s, as expected from our previous study, and very similar Na kinetics were observed when cytoplasmic Na changes were inferred from decay of Na channel currents after rapid Na application and removal (Fig. 8B). Again, the estimated time constant for Na turnover to the pipette was 22 s. One potential objection to our detailed interpretation is the assumption of a mixing volume for cytoplasmic Na of only 33% of total myocyte volume. This is consistent with the known volume of mitochondria in murine myocytes (e.g. (Swietach et al., 2015)) together with evidence that hydration water of myocyte myofilaments amounts to ∼20% of total cellular water (Drewnowska and Baumgarten, 1991; Rorschach et al., 1991).

We can only speculate why repeated depolarizations might activate Na/K pump currents for prolonged times (Skogestad et al., 2019). Without knowing Na channel reversal potentials, it is possible that Na/K pumps are stimulated by a regulatory mechanism activated by rapid depolarizations. Control of Ca is one potential concern, since Ca elevations stimulate very markedly Na/K pump currents in murine myocyte (Lu et al., 2016). To account for Na turnover, we must assume that the cytoplasmic D_Na_ in murine myocytes is at least 5 x 10^-6^ cm^2^/s, not more that 2.5-fold reduced from diffusion in free water.

### Nucleotide diffusion in cardiac myocytes

We have attempted to define the diffusivity of MgATP in murine cardiac cytoplasm in four ways. First we determined the time courses of Na/K pump inhibition and recovery upon perfusing AMPPNP into and out of the patch pipette tip (Fig. 9A). Second, we determined the time course of pump current decline during patch clamp with ATP-free pipette solutions (Fig. 9B). Third, we determined the time course with which anions, including MgATP, diffuse into myocytes from the patch pipette with Na as the coupled monovalent cation (Figs.10 and 11). Fourth, we compared diffusion of NaCl and Na(Mg)ATP into sarcolemma-permeabilized myocytes constrained within large-diameter pipette tips (Fig. 12A). In the first three protocols, the results indicate that MgATP diffuses with a coefficient of 0.2 to 0.5 x 10^-7^ cm^2^/s. These values are roughly three-fold smaller than coefficients determined for skeletal muscle (see Introduction) and 8 to 10 times smaller than for ATP diffusion in free water. In the experiments employing permeabilized myocytes, MgATP diffuses 11 times slower than Cl, which equates to about four-times slower than in free water. All results are consistent with cardiac myofilaments and mitochondrial networks restricting diffusion more than skeletal myofilaments and mitochondria. As a result, local metabolite gradients may be greater and occur in a more structured manner than in skeletal muscle. The reasons for differences observed in cardiac versus skeletal myofilaments (Fig. 13), and also between mouse strains, are a matter of speculation. One suspicion is that the greater passive stiffness of cardiac muscle reflects different packing arrangements of filaments that could in turn influence diffusion within cardiac myofilaments. An involvement of cardiac titins is one speculative possibility (LeWinter and Granzier, 2010) that might give rise to diffusion restrictions that depend on titin oxidation status (Loescher et al., 2020).

### Diffusion of fluorophores in cardiac myocytes

That commonly employed fluorophores diffuse 15 to 50 times slower than in free water explains why cardiac electrophysiologists typically employ AM esters to load fluorophores, even when employing patch clamp. Among the different fluorophores examined, only sulforhodamine binds profusely to cytoplasmic constituents, while FITC-PEGs show essentially no evidence of binding. Notably, a 10kD PEG diffuses faster than a 2kD PEG in myoplasm, and others have also noted examples in which 10 kD solutes diffuse faster than 1kD molecules in myocytes (Illaste et al., 2012). One possible explanation is that larger fluorophores are not moving via simple diffusion but are moving by other mechanisms that involve cytoskeleton and/or conformational changes of myofilaments.

### Implications of dual diffusion restrictions in cardiac myocytes

Diffusion restrictions in murine myocytes are much greater than described for molecular crowding in bacteria (McGuffee and Elcock, 2010), as well as in non-muscle mammalian cells (Verkman, 2002; Rivas and Minton, 2016). Particularly for bacteria, diffusion restrictions are characterized in much detail (McGuffee and Elcock, 2010; Gallet et al., 2017; Bohrer and Xiao, 2020), and molecular simulations are quite advanced (Trovato and Tozzini, 2014; Bohrer and Xiao, 2020). In those models, only a two to three fold reduction of diffusion coefficients occurs as hydrated radii increase from 1 to 4 nm (i.e. as molecular weight increases to >20 kD). This is similar to results for mammalian cells that do not contain a dense contractile protein network or extensive mitochondrial networks (Verkman, 2002). Gel-like protein meshes clearly restrict diffusion in more complex ways than molecular crowding by diffusible macromolecules (e.g. (Lauffer, 1961; Johnson et al., 1996; Labille et al., 2007)). Diffusion of fluorophores in agarose is only mildly restricted (Shoga et al., 2017), and the diffusion of small solutes in cartilage is restricted by only about two-fold (Burstein et al., 1993). The obvious structural similarity between cardiac myoplasm and gelatins is they are both protein networks with cross-linkages, and we verify in Sup. Fig. S9 that diffusion restrictions in gelatins are essentially lost upon warming just enough to disengage gel linkages.

As outlined in the Introduction, it has been repeatedly suggested that mitochondrial networks restrict diffusion in cardiac myocytes, and reduced diffusivity of ATP and carboxyfluorescein in VDAC1 knockout myocytes (Fig. 15) provides strong new evidence for these suggestions. It has also been suggested that the sarcoplasmic reticulum can restrict diffusion in myocytes by effectively surrounding myofilaments (Fabiato, 1985). Although the geometric prerequisites for diffusion restriction by cytoplasmic organelles are not rigorously established, mathematical models have been developed that appear to predict the basic outcomes of the present study (Ramay and Vendelin, 2009). As outlined in Figs. 13 to 16, our data suggest that diffusion restrictions arise approximately equally from cardiac myofilaments and mitochondrial networks. As a result, porin channels of the outer mitochondrial membrane may facilitate diffusion by allowing solute passage through mitochondrial intermembrane spaces. Given that mitochondria make up 35% of the murine myocyte volume (Dedkova and Blatter, 2012) and that the free mixing volume of the myofilament space is likely not more than 30% of myocyte volume (Lu and Hilgemann, 2017), the undulating 8 nm intermembrane space of mitochondria will likely constitute several percent of the free cellular mixing volume. Accordingly, a 5- to 10- fold restriction of solute diffusion in myofilament spaces can plausibly lend a preference to solute passage through intermembrane spaces. Of course, it is prerequisite that diffusion in the mitochondrial intermembrane spaces takes place without significant restriction.

In conclusion, long-distance diffusion coefficients for small solutes decrease profoundly over the molecular weight range of 200 to 1000 in the cytoplasm of BL6 murine cardiac myocytes. Fluorophores are restricted by 20 to 50-fold, while ATP appears to diffuse 8- to 10-times slower than in free water. Further work is required to establish how and if these restrictions affect specific myocyte functions and whether cardiac diffusion restrictions are subject to regulatory influences (i.e. by regulating porin channel properties and/or the stiffness of cardiac myofilaments). It remains an open question whether protein networks (e.g. actin cytoskeleton) can restrict and/or channel solute diffusion to a physiologically relevant extent in submicroscopic spaces next to membranes and membrane proteins (Chu et al., 2012).

## Acknowledgements

We thank Fang-Min Lu for technical help and Orson Moe (UTSouthwestern) for support, discussion, and encouragement. Supported by NIH HL119843.

## Reference List

Agarwal, S.R., C.E. Clancy, and R.D. Harvey. 2016. Mechanisms Restricting Diffusion of Intracellular cAMP. Scientific reports. 6:19577.

Anflous, K., D.D. Armstrong, and W.J. Craigen. 2001. Altered mitochondrial sensitivity for ADP and maintenance of creatine-stimulated respiration in oxidative striated muscles from VDAC1-deficient mice. J Biol Chem. 276:1954–1960.

Arrio-Dupont, M., S. Cribier, G. Foucault, P.F. Devaux, and A. d’Albis. 1996. Diffusion of fluorescently labeled macromolecules in cultured muscle cells. Biophys J. 70:2327–2332.

Artemov, V.G.V., A.,A.;Sysoev, N.N.; Volkov, A.A. 2015. Conductivity of aqueous HCl, NaOH and NaCl solutions: Is water just a substrate? EPL. 109:1–7.

Baylor, S.M., and S. Hollingworth. 1998. Model of sarcomeric Ca2+ movements, including ATP Ca2+ binding and diffusion, during activation of frog skeletal muscle. The Journal of general physiology. 112:297–316.

Beaugé, L.A., and I.M. Glynn. 1980. The equilibrium between different conformations of the unphosphorylated sodium pump: effects of ATP and of potassium ions, and their relevance to potassium transport. The Journal of physiology. 299:367–383.

Blatter, L.A., and W.G. Wier. 1990. Intracellular diffusion, binding, and compartmentalization of the fluorescent calcium indicators indo-1 and fura-2. Biophysical journal. 58:1491–1499.

Bohrer, C.H., and J. Xiao. 2020. Complex Diffusion in Bacteria. Advances in experimental medicine and biology. 1267:15–43.

Bowen, W.J., and H.L. Martin. 1963. A STUDY OF DIFFUSION OF ATP THROUGH GLYCEROL-TREATED MUSCLE. Archives of biochemistry and biophysics. 102:286–292.

Bowen, W.J., and H.L. Martin. 1964. THE DIFFUSION OF ADENOSINE TRIPHOSPHATE THROUGH AQUEOUS SOLUTIONS. Archives of biochemistry and biophysics. 107:30–36.

Burstein, D., M.L. Gray, A.L. Hartman, R. Gipe, and B.D. Foy. 1993. Diffusion of small solutes in cartilage as measured by nuclear magnetic resonance (NMR) spectroscopy and imaging. Journal of orthopaedic research : official publication of the Orthopaedic Research Society. 11:465–478.

Carmeliet, E. 1992. A fuzzy subsarcolemmal space for intracellular Na+ in cardiac cells? Cardiovascular research. 26:433–442.

Chen, C., T. Nakamura, and Y. Koutalos. 1999. Cyclic AMP diffusion coefficient in frog olfactory cilia. Biophysical journal. 76:2861–2867.

Chu, H., E. Puchulu-Campanella, J.A. Galan, W.A. Tao, P.S. Low, and J.F. Hoffman. 2012. Identification of cytoskeletal elements enclosing the ATP pools that fuel human red blood cell membrane cation pumps. Proc Natl Acad Sci U S A. 109:12794–12799.

Collins, A., A.V. Somlyo, and D.W. Hilgemann. 1992. The giant cardiac membrane patch method: stimulation of outward Na(+)-Ca2+ exchange current by MgATP. J Physiol. 454:27–57.

de Graaf, R.A., A. van Kranenburg, and K. Nicolay. 2000. In vivo (31)P-NMR diffusion spectroscopy of ATP and phosphocreatine in rat skeletal muscle. Biophysical journal. 78:1657–1664.

Dedkova, E.N., and L.A. Blatter. 2012. Measuring mitochondrial function in intact cardiac myocytes. J Mol Cell Cardiol. 52:48–61.

Drewnowska, K., and C.M. Baumgarten. 1991. Regulation of cellular volume in rabbit ventricular myocytes: bumetanide, chlorothiazide, and ouabain. The American journal of physiology. 260:C122–131.

Fabiato, A. 1985. Rapid ionic modifications during the aequorin-detected calcium transient in a skinned canine cardiac Purkinje cell. J Gen Physiol. 85:189–246.

Friedman, L.K., E.O. 1930. The structure of gelatin gels from studies of diffusion. The Journal of the American Chemical Society. 52:1295–1304.

Gallet, R., C. Violle, N. Fromin, R. Jabbour-Zahab, B.J. Enquist, and T. Lenormand. 2017. The evolution of bacterial cell size: the internal diffusion-constraint hypothesis. The ISME journal. 11:1559–1568.

Garcia, A., C.C. Liu, F. Cornelius, R.J. Clarke, and H.H. Rasmussen. 2016. Glutathionylation-Dependence of Na(+)-K(+)-Pump Currents Can Mimic Reduced Subsarcolemmal Na(+) Diffusion. Biophysical journal. 110:1099–1109.

Gendron, P.O., F. Avaltroni, and K.J. Wilkinson. 2008. Diffusion coefficients of several rhodamine derivatives as determined by pulsed field gradient-nuclear magnetic resonance and fluorescence correlation spectroscopy. J Fluoresc. 18:1093–1101.

Glonek, T. 1992. 31P NMR of Mg-ATP in dilute solutions: complexation and exchange. Int J Biochem. 24:1533–1559.

Gribbon, P., and T.E. Hardingham. 1998. Macromolecular diffusion of biological polymers measured by confocal fluorescence recovery after photobleaching. Biophysical journal. 75:1032–1039.

Hasselbach, W. 1952. Die Diffusionskonstante des Adenosinteriphosphats im Inneren der Muskelfaser. NZeitschrift fuer Naturforschung B. 7:334–338.

Hayes, J.S., and L.L. Brunton. 1982. Functional compartments in cyclic nucleotide action. Journal of cyclic nucleotide research. 8:1–16.

Hilgemann, D.W., and C.C. Lu. 1998. Giant membrane patches: improvements and applications. Methods Enzymol. 293:267–280.

Hubley, M.J., R.C. Rosanske, and T.S. Moerland. 1995. Diffusion coefficients of ATP and creatine phosphate in isolated muscle: pulsed gradient 31P NMR of small biological samples. NMR in biomedicine. 8:72–78.

Iamshanova, O., P. Mariot, V. Lehen’kyi, and N. Prevarskaya. 2016. Comparison of fluorescence probes for intracellular sodium imaging in prostate cancer cell lines. European biophysics journal : EBJ. 45:765–777.

Illaste, A., M. Laasmaa, P. Peterson, and M. Vendelin. 2012. Analysis of molecular movement reveals latticelike obstructions to diffusion in heart muscle cells. Biophysical journal. 102:739–748.

Jepihhina, N., N. Beraud, M. Sepp, R. Birkedal, and M. Vendelin. 2011. Permeabilized rat cardiomyocyte response demonstrates intracellular origin of diffusion obstacles. Biophysical journal. 101:2112–2121.

Johnson, E.M., D.A. Berk, R.K. Jain, and W.M. Deen. 1996. Hindered diffusion in agarose gels: test of effective medium model. Biophysical journal. 70:1017–1023.

Jones, D.P. 1986. Intracellular diffusion gradients of O2 and ATP. The American journal of physiology. 250:C663–675.

Jurevicius, J., and R. Fischmeister. 1996. cAMP compartmentation is responsible for a local activation of cardiac Ca2+ channels by beta-adrenergic agonists. Proceedings of the National Academy of Sciences of the United States of America. 93:295–299.

Kamceva, J.S., R.; Janga, E-S.; Yana, N.; Moeb, N.; Paula, D.P.; Freemana, B.D. 2018. Salt concentration dependence of ionic conductivity in ion exchange membranes. Journal of Membrane Science. 547:123–133.

Kim, J., R. Gupta, L.P. Blanco, S. Yang, A. Shteinfer-Kuzmine, K. Wang, J. Zhu, H.E. Yoon, X. Wang, M. Kerkhofs, H. Kang, A.L. Brown, S.J. Park, X. Xu, E. Zandee van Rilland, M.K. Kim, J.I. Cohen, M.J. Kaplan, V. Shoshan-Barmatz, and J.H. Chung. 2019. VDAC oligomers form mitochondrial pores to release mtDNA fragments and promote lupus-like disease. Science (New York, N.Y.). 366:1531–1536.

Konishi, M., and M. Watanabe. 1995. Molecular size-dependent leakage of intracellular molecules from frog skeletal muscle fibers permeabilized with beta-escin. Pflugers Arch. 429:598–600.

Kramer, E.M., N.L. Frazer, and T.I. Baskin. 2007. Measurement of diffusion within the cell wall in living roots of Arabidopsis thaliana. J Exp Bot. 58:3005–3015.

Kushmerick, M.J., and R.J. Podolsky. 1969. Ionic mobility in muscle cells. Science (New York, N.Y.). 166:1297–1298.

Labille, J., N. Fatin-Rouge, and J. Buffle. 2007. Local and average diffusion of nanosolutes in agarose gel: the effect of the gel/solution interface structure. Langmuir : the ACS journal of surfaces and colloids. 23:2083–2090.

Lauffer, M.A. 1961. Theory of diffusion in gels. Biophysical journal. 1:205–213.

Lee, S.H. 2020. Dynamic and Static Properties of Aqueous NaCl Solutions at25°C as a Function of NaCl Concentration:A Molecular Dynamics Simulation Study. Journal of Chemistry. 2020.

LeWinter, M.M., and H. Granzier. 2010. Cardiac titin: a multifunctional giant. Circulation. 121:2137–2145.

Loescher, C.M., M. Breitkreuz, Y. Li, A. Nickel, A. Unger, A. Dietl, A. Schmidt, B.A. Mohamed, S. Kötter, J.P. Schmitt, M. Krüger, M. Krüger, K. Toischer, C. Maack, L.I. Leichert, N. Hamdani, and W.A. Linke. 2020. Regulation of titin-based cardiac stiffness by unfolded domain oxidation (UnDOx). Proc Natl Acad Sci U S A. 117:24545–24556.

Lu, F.M., C. Deisl, and D.W. Hilgemann. 2016. Profound regulation of Na/K pump activity by transient elevations of cytoplasmic calcium in murine cardiac myocytes. eLife. 5.

Lu, F.M., and D.W. Hilgemann. 2017. Na/K pump inactivation, subsarcolemmal Na measurements, and cytoplasmic ion turnover kinetics contradict restricted Na spaces in murine cardiac myocytes. J Gen Physiol. 149:727–749.

Masaro, L.Z., X.,X. 1999. Physical models of diffusion for polymer solutions, gels and solids. Prog. Polym. Sci. 24:731–775.

McGuffee, S.R., and A.H. Elcock. 2010. Diffusion, crowding & protein stability in a dynamic molecular model of the bacterial cytoplasm. PLoS computational biology. 6:e1000694.

Miller, D.S., and S.B. Horowitz. 1986. Intracellular compartmentalization of adenosine triphosphate. J Biol Chem. 261:13911–13915.

Naumova, A.V., R.G. Weiss, and V.P. Chacko. 2003. Regulation of murine myocardial energy metabolism during adrenergic stress studied by in vivo 31P NMR spectroscopy. Am J Physiol Heart Circ Physiol. 285:H1976–1979.

Palmer, R.E., A.J. Brady, and K.P. Roos. 1996. Mechanical measurements from isolated cardiac myocytes using a pipette attachment system. The American journal of physiology. 270:C697–704.

Pape, P.C., D.S. Jong, W.K. Chandler, and S.M. Baylor. 1993. Effect of fura-2 on action potential-stimulated calcium release in cut twitch fibers from frog muscle. The Journal of general physiology. 102:295–332.

Piquereau, J., F. Caffin, M. Novotova, C. Lemaire, V. Veksler, A. Garnier, R. Ventura-Clapier, and F. Joubert. 2013. Mitochondrial dynamics in the adult cardiomyocytes: which roles for a highly specialized cell? Front Physiol. 4:102.

Popel, A.S. 1989. Theory of oxygen transport to tissue. Critical reviews in biomedical engineering. 17:257–321.

Ramay, H.R., and M. Vendelin. 2009. Diffusion restrictions surrounding mitochondria: a mathematical model of heart muscle fibers. Biophys J. 97:443–452.

Richards, M., O. Lomas, K. Jalink, K.L. Ford, R.D. Vaughan-Jones, K. Lefkimmiatis, and P. Swietach. 2016. Intracellular tortuosity underlies slow cAMP diffusion in adult ventricular myocytes. Cardiovascular research. 110:395–407.

Rivas, G., and A.P. Minton. 2016. Macromolecular Crowding In Vitro, In Vivo, and In Between. Trends in biochemical sciences. 41:970–981.

Robinson, J.D. 1980. Binding to the high-affinity substrate site of the (Na+ + K+)-dependent ATPase. Journal of bioenergetics and biomembranes. 12:165–174.

Rorschach, H.E., C. Lin, and C.F. Hazlewood. 1991. Diffusion of water in biological tissues. Scanning microscopy. Supplement. 5:S1–9; discussion S9-10.

Saks, V.A., Z.A. Khuchua, E.V. Vasilyeva, O. Belikova, and A.V. Kuznetsov. 1994. Metabolic compartmentation and substrate channelling in muscle cells. Role of coupled creatine kinases in in vivo regulation of cellular respiration--a synthesis. Molecular and cellular biochemistry. 133–134:155–192.

Sanabria, H., Y. Kubota, and M.N. Waxham. 2007. Multiple diffusion mechanisms due to nanostructuring in crowded environments. Biophysical journal. 92:313–322.

Shoga, J.S., B.T. Graham, L. Wang, and C. Price. 2017. Direct Quantification of Solute Diffusivity in Agarose and Articular Cartilage Using Correlation Spectroscopy. Annals of biomedical engineering. 45:2461–2474.

Sidell, B.D., and J.R. Hazel. 1987. Temperature affects the diffusion of small molecules through cytosol of fish muscle. The Journal of experimental biology. 129:191–203.

Simson, P., N. Jepihhina, M. Laasmaa, P. Peterson, R. Birkedal, and M. Vendelin. 2016. Restricted ADP movement in cardiomyocytes: Cytosolic diffusion obstacles are complemented with a small number of open mitochondrial voltage-dependent anion channels. Journal of molecular and cellular cardiology. 97:197–203.

Skogestad, J., G.T. Lines, W.E. Louch, O.M. Sejersted, I. Sjaastad, and J.M. Aronsen. 2019. Evidence for heterogeneous subsarcolemmal Na(+) levels in rat ventricular myocytes. American journal of physiology. Heart and circulatory physiology. 316:H941–H957.

Song, Z.S., A.; Eady, J.; Zhao, H.; Olubajo, O. 2008. Determining nucleotide acidity and cation binding constans by 31P NMR. Canadian Journal of Analytical Sciences and Spectroscopy. 53:45–51.

Stockbridge, R.B., and R. Wolfenden. 2009. The intrinsic reactivity of ATP and the catalytic proficiencies of kinases acting on glucose, N-acetylgalactosamine, and homoserine: a thermodynamic analysis. J Biol Chem. 284:22747–22757.

Swietach, P., K.W. Spitzer, and R.D. Vaughan-Jones. 2015. Na⁺ ions as spatial intracellular messengers for co-ordinating Ca²⁺ signals during pH heterogeneity in cardiomyocytes. Cardiovascular research. 105:171–181.

Timmerman, M.P., and C.C. Ashley. 1986. Fura-2 diffusion and its use as an indicator of transient free calcium changes in single striated muscle cells. FEBS letters. 209:1–8.

Trovato, F., and V. Tozzini. 2014. Diffusion within the cytoplasm: a mesoscale model of interacting macromolecules. Biophysical journal. 107:2579–2591.

Verdonck, F., K. Mubagwa, and K.R. Sipido. 2004. [Na(+)] in the subsarcolemmal ‘fuzzy’ space and modulation of [Ca(2+)](i) and contraction in cardiac myocytes. Cell Calcium. 35:603–612.

Verkman, A.S. 2002. Solute and macromolecule diffusion in cellular aqueous compartments. Trends Biochem Sci. 27:27–33.

Waggoner, R.A.B., F.D.; Lang, J.C. 1995. Diffusion in Aqueous Solutions of Poly(ethylene glycol) at Low Concentrations. Macromolecules. 28:2658–2664.

Wang, T.M., and D.W. Hilgemann. 2008. Ca-dependent nonsecretory vesicle fusion in a secretory cell. J Gen Physiol. 132:51–65.

Weeber, E.J., M. Levy, M.J. Sampson, K. Anflous, D.L. Armstrong, S.E. Brown, J.D. Sweatt, and W.J. Craigen. 2002. The role of mitochondrial porins and the permeability transition pore in learning and synaptic plasticity. J Biol Chem. 277:18891–18897.

Weiss, J.N., and S.T. Lamp. 1987. Glycolysis preferentially inhibits ATP-sensitive K+ channels in isolated guinea pig cardiac myocytes. *Science (New York*, N.Y.). 238:67–69.

Weiss, J.N., and S.T. Lamp. 1989. Cardiac ATP-sensitive K+ channels. Evidence for preferential regulation by glycolysis. The Journal of general physiology. 94:911–935.

Wheatley, D.N. 2003. Diffusion, perfusion and the exclusion principles in the structural and functional organization of the living cell: reappraisal of the properties of the ‘ground substance’. The Journal of experimental biology. 206:1955–1961.

Widodo, C.S.S., H; Santosa, D.R. 2018. The effect of NaCl concentration on the ionic NaCl solutions electrical impedance value using electrochemical impedance spectroscopy methods. AIP Conference Proceedings. 050003.

Zong, X.G., M. Dugas, and P. Honerjäger. 1992. Relation between veratridine reaction dynamics and macroscopic Na current in single cardiac cells. The Journal of general physiology. 99:683–697.

