## Supplemental Data for "Pore-like diffusion barriers in murine cardiac myocytes"

**Description of Supplementary Materials**. *First,* we provide supplementary data concerning cardiac myocytes employed in this study. This included further examples of fluorophore diffusion in myocytes, a description of experiments with Na dyes to attempt to monitor Na changes in myocytes, and the presentation of Western blots that document VDAC1 knockout and lack of compensation of VDAC1 by other VDAC isoforms in the VDAC1 knockout myocytes. *Second,* we provide an analysis of diffusion in gelatins that has relevance to diffusion in myocytes. We demonstrate with both optical and electrical methods that diffusion in gelatins is restricted with a strong dependence on molecular weight, similar to diffusion in myocytes. Similar to diffusion in myocytes, ATP diffusion is restricted by nearly 10-fold in 10 gram percent gelatin, and restrictions are effectively lost when the gelling state is lost by warming to 36°C.

### 1. Supplementary materials concerning cardiac myocytes.

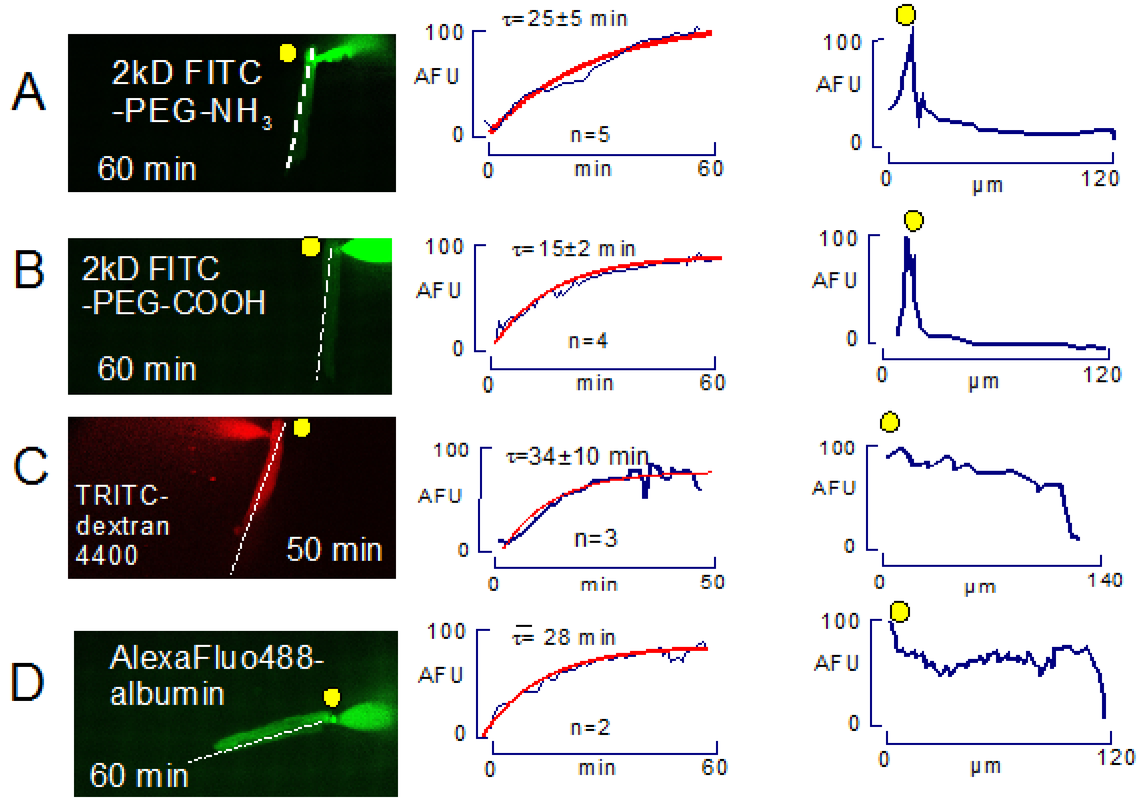


**Figure S1. Diffusion of four additional fluorescent molecules into the cytoplasm of patch clamped murine myocytes.** Same experimental conditions as Figs. 6 and 7. **A.** 2kD FITC-labelled PEG with one amino group (100 μM). **B**. 2kD FITC-labelled PEG with one carboxyl group (100 μM). **C.** 4.4 kD TRITC-labelled dextran (10 μM). **D**. AlexaFluo488-labelled albumin (100 μM). Time constants determined for the positively and negatively charged PEGs were not significantly different. The 4 kD dextran equilibrates significantly slower, but a fluorescent albumin diffuses quite rapidly. Similar to results for GFP (Fig.7) the fluorescent albumin did not show distinct gradients and myocyte fluorescence remains substantially lower than pipette fluorescence.

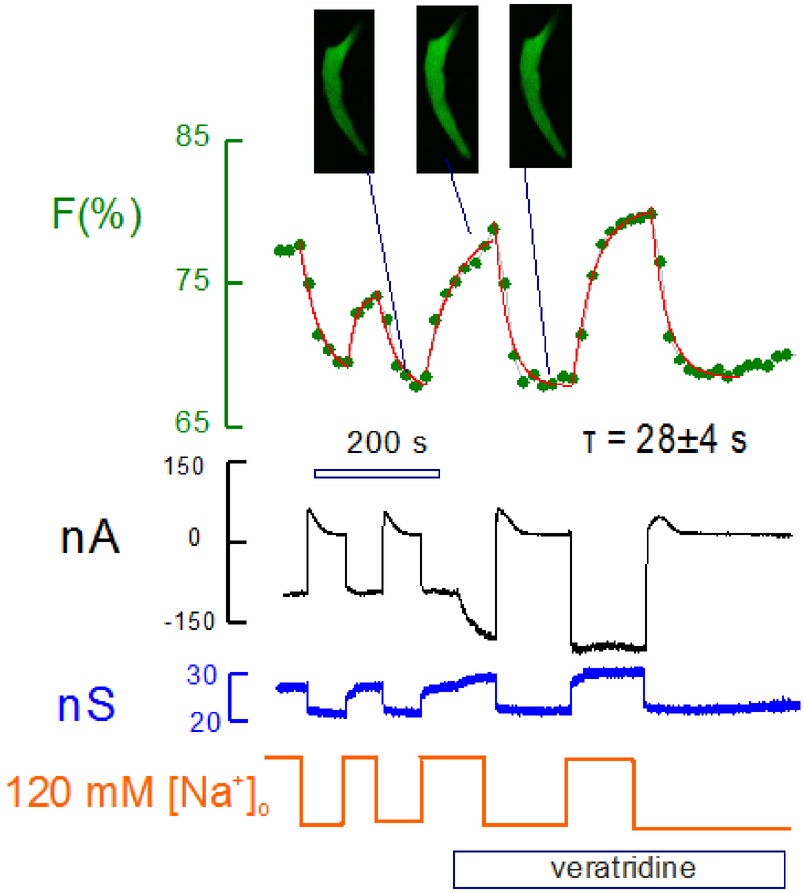


**Figure S2.** Optical measurement of Na concentration changes during and after Na influx activated by veratridine (3 µM). Myocytes were loaded with CoronaGreen AM Na-selective dye (5 µM AM) for 30 min at 25°C (Iamshanova et al., 2016). Myocytes were patch clamped using Na-free NMG solutions as in Fig. 8. Na was applied and removed multiple times without veratridine and subsequently with veratridine. In the presence of extracellular Na (120 mM), cytoplasmic CoronaGreen fluorescence increases by ~5% and decreases again upon removal of extracellular Na. Fluorescence changes increase by about a factor of 2 in the presence of veratridine. Each fluorescence change was fit to falling or rising asymptotic exponential functions (red curves), and the average time constant amounted to 28 s. The magnitudes of luorescence changes (5 to 10%) are consistent with Na changes of 2 to 4 mM, assuming a $K_d$ of 10 mM for Na binding by CoronaGreen.

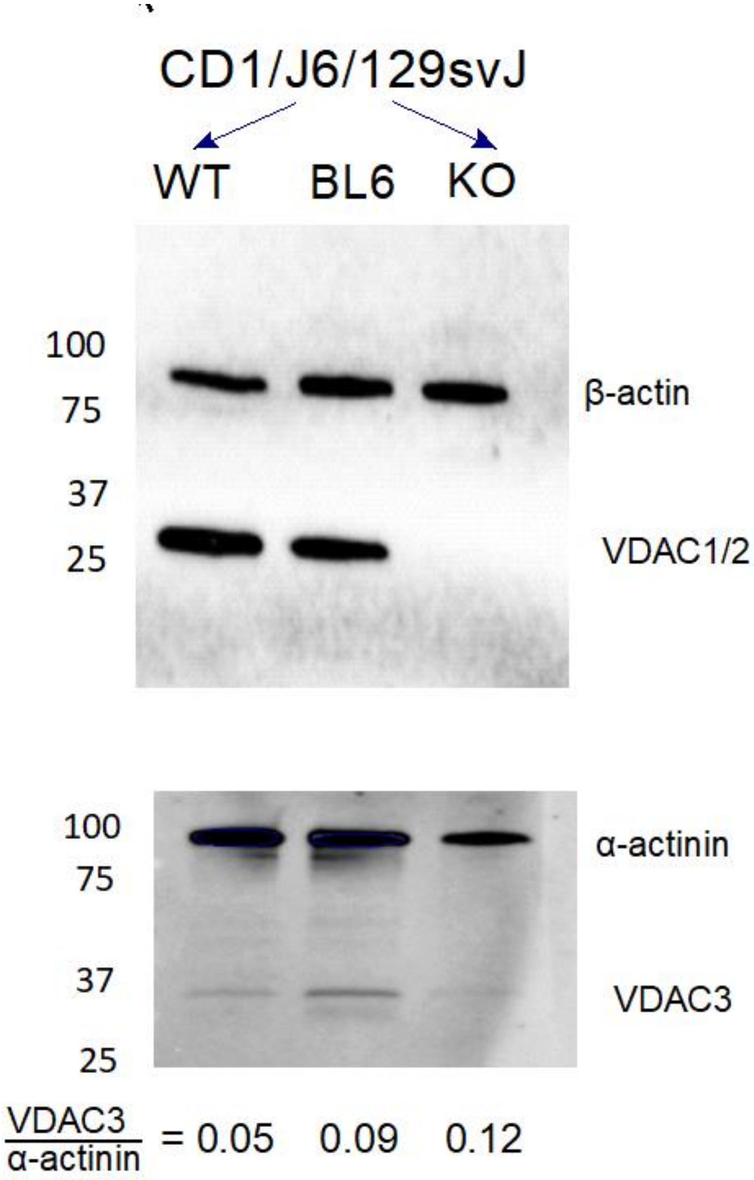


**Figure S3.** Western blots of VDAC isoforms in WT ‘CD1’ cardiac myocytes, WT BL6 myocytes, and VDAC1-deficient ‘CD1’ myocytes. Antibodies employed were VDAC1/2 antibody 10866-1-AP and VDAC3 55260-1-AP from Proteintech. VDAC expression is similar in the two mouse strains, and the knockout mice to not express either VDAC1 or VDAC2.

#### 2. Supplementary materials concerning diffusion in gelatin.

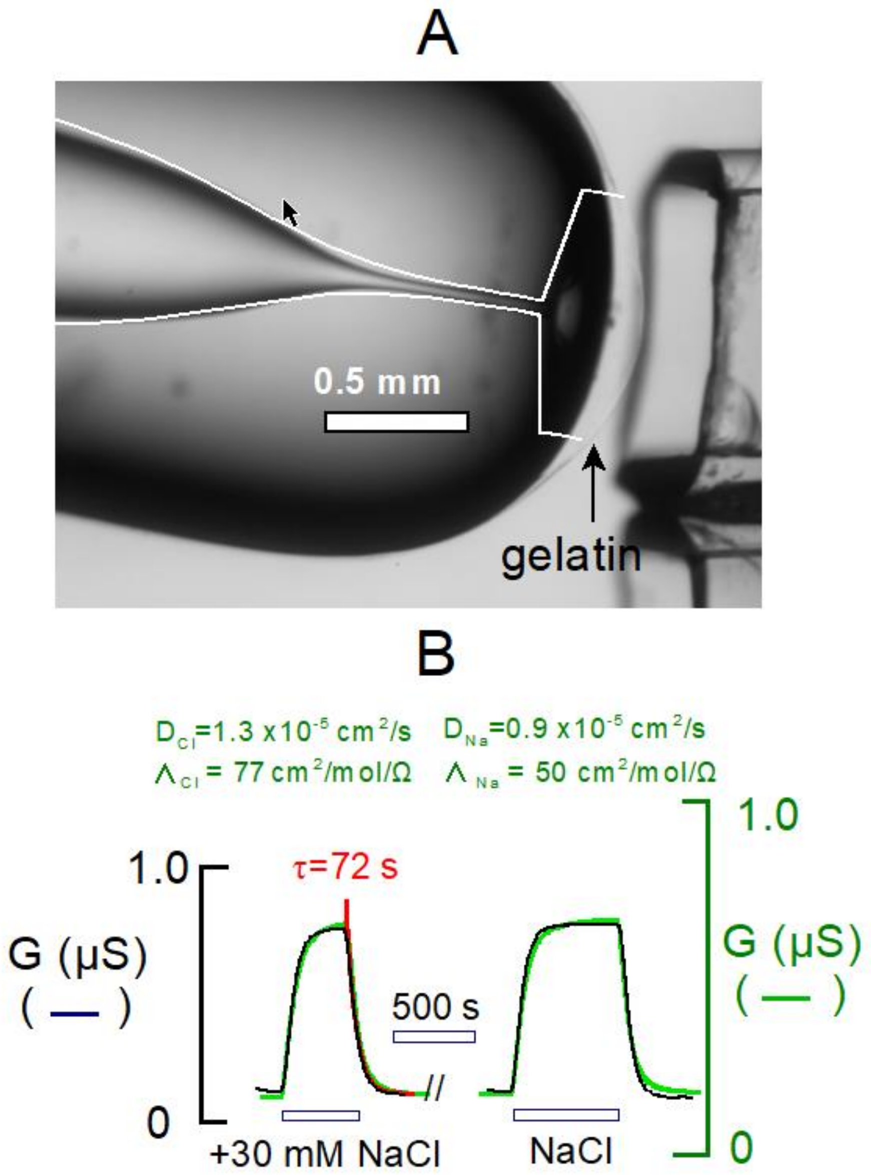


**Figure S4.** Conductance measurements of diffusion in gelatin. **A.** Borosilicate glass pipettes with cylindrical orifices were formed on a glass flame. Tips were filled over ~ 6 mm with warm 10 gram% gelatin solution and backfilled with low ionic strength solution. A 150 to 250 micron thick gelatin coat adhered to the tip. Sinusoidal voltage oscillations were applied to monitor conductance of the tip. **B.** Typical conductance records for addition and removal of 30 mM NaCl. Diffusion coefficients were estimated via simulations (green) with approximated inner tip dimensions (white lines in A). Optimal diffusion coefficients for Na and Cl, adjusted in a ratio of 1.5:1, were 33% less than those for free water (Robinson, 1965). Reasonable estimates of coupled diffusion coefficients can also be made from exponential fits of the decaying conductance curves, assuming a median diffusion distance of 0.35 mm (see red line in left record).

**Additional information.** The tips of borosilicate glass pipettes 2 mm in diameter were melted on a gas flame until the openings collapsed to form nearly cylindrical vestibules ~40 µm in diameter and 0.25 to 0.5 mm in length, as shown in Fig. S4A. Distal to the vestibule, the inner diameter increased to >300 µm over a distance of ~0.35 mm (not clearly visible in Fig.S4A). Gelatin (10% wt/volume) was prepared in a low ionic strength solution designated as 'background solution' (3 mM KCl, 0.5 mM HEPES, and 0.25 mM EGTA, set to pH 7.2 with KOH). The pipette tip was dipped into the warm, liquid gelatin solution until the gelatin filled the tip to a height of approximately 4 mm, and the tip was back-filled with the low ionic strength solution. As evident in Fig. S4A, gelatin coated the outer pipette surface at a thickness of about 150 µm. Sinusoidal voltage oscillations were applied (0.2 to 1 kHz, 2 to 6 mV) via an Axopatch 1C patch clamp, employed in 'tracking' mode to null the mean applied current with a time constant of ~2s. The tip conductance was then monitored over time using our own software (Wang and Hilgemann, 2008). After equilibration of the gelatin-filled tip with background solution, solutions with higher ionic strengths were applied to the outside of the pipette. Usually, solutions were applied through 1mm square pipettes via gravity-feed, as in patch clamp experiments with myocytes. Alternatively, for compounds that were relatively precious, the pipette tip was inserted into small vials containing 200 µL of solutions, connected to one another and to the patch clamp ground via chlorinated silver wires. The potentials which developed upon changing ionic strength reflected accurately the Cl concentrations of solutions, and they were negated effectively within 3 seconds by the 'tracking' mode of the patch clamp.

For the simulations, the pipette dimensions were reconstructed digitally (white lines in Fig. S4A) and used to simulate diffusion through the pipette tip. Gelatin coating of the pipette tip was simulated as a 0.26 mm thick column with a 0.5 mm radius. Then, the diffusion coefficients for

Na and Cl were varied with the ratio of Cl to Na diffusion coefficients fixed at the ratio found in free water, 1.54 (Robinson, 1965). Optimal conductance kinetics were generated with diffusion coefficients (1.3 and 0.9 x $10^{-5}$ $cm^2/s$ for Cl and Na respectively). When conductances were calculated with equivalent conductivities for Na and Cl found in free water (Robinson, 1965), the calculated conductances (green line in Fig. S4B) were about 25% less than measured conductances (see green scale bar in Fig. S4B). We conclude that NaCl diffusion is restricted by not more than 40% by 10 gram% gelatin. We mention that optimal simulation parameters often generated a rising phase that was slightly too slow with a falling phase that was slightly too fast. We also point out that the falling conductance curves can be fit accurately over their terminal two-thirds, roughly, with an exponential function (red curve in the left record). Simulations described in Methods verify that coupled diffusion coefficients, $(2 \times D_{Na} \times D_{Cl})/(D_{Na} + D_{Cl})$ in this case, can be accurately estimated from the average diffusion distance (~0.69 x 0.6 cm) and the time constant of conductance decay with $D=x^2/2\tau$. In the present case, our estimate , 1.2 x $10^{-5}$ $cm^2/s$, is very close to the estimate derived by simulation.

The simulations were carried out in Matlab (MatWorks) using essentially the same routines described previously (Wang and Hilgemann, 2008; Swietach et al., 2015) with accuracy checking to within line-width of graphs provided. When appropriate, the shape of pipette tips was approximated in microns by a modified Hill equation. For the example in Fig. S4B,

(S1) radius in µm $= 480 \times L^4/(L^4+1500^4)+30$ ,

where L is the distance from the pipette orifice in microns. For Fig. S4B, we simulated the gelatin coating of the pipette tip as a disk with a radius of 0.5 mm and a width of 0.22 mm. When we compared simulations using this approach to simpler simulations of diffusion through a cylinder of appropriate length (0.3 to 0.5 mm) with fixed ion concentrations at opposite ends, the optimal diffusion coefficients derived were not more than 50% different. This reflects the fact that pipette orifices are approximately straight over 0.4 mm with pore diameters increasing quite steeply thereafter.

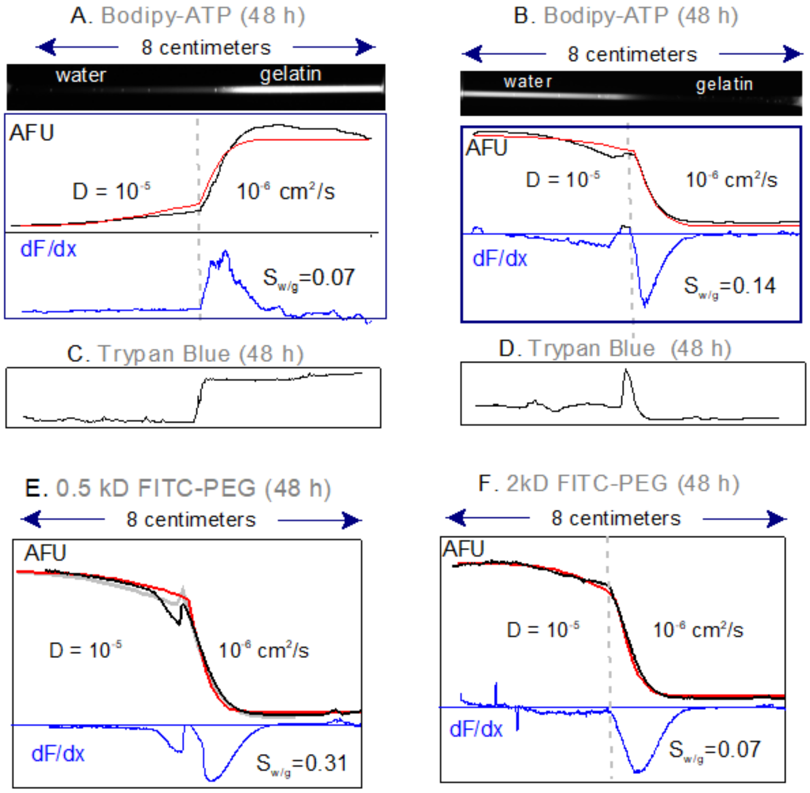


**Figure S5.** Fluorophore diffusion in 8 cm glass pipettes filled one-half with 10% gelatin. Simulations (red curves) were used to determine best estimates of diffusion constants given in the figure **A, B.** Gelatin solution with and without Bodipy-FL-ATP (5 µM) was loaded into one-half of 2 pipettes each. Gelatin-free solution without and with Bodipy-FL-ATP was loaded into the other pipette halves, respectively. Fluoresence profiles (black) and their derivatives (blue) are shown after 48 h for dye diffusion out of (A) and into (B) gelatin. Without gelatin, diffusion progressed over the entire pipette distance. The ratio of the peak dF/dx in water to that in gelatin ($S_{w/g}$) was 0.07 and 0.14, indicating that diffusion was ~10-fold restricted in gelatin **C, D.** Equivalent experiments with Trypan Blue (30 µM), which fluoresces when bound to proteins and other cell constituents. Over 48 h Trypan Blue effectively does not diffuse out of gelatin (C), nor does it diffuse from gelatin-free solution into gelatin (D). **E,F.** Diffusion of 0.5 kD and 2kD PEG from gelatin-free solution into gelatin, respectively. Based on the ratio of slopes ($S_{w/g}$), the 0.5 kD PEG is 3.3 fold restricted in gelatin and the 2kD PEG is 14-fold restricted. Results were similar in 3 further experiments.

| Observed solute | $S_{w/g}$ |
|---|---|
| Bodipy-FL-ATP | 0.07 |
| 0.5 kD FITC-PEG | 0.31 |
| 2 kD FITC-PEG | 0.07 |
| carboxyfluorescein | 0.06 |
| sulforhodamine | 0.16 |
| ASG2 | 0.12 |
| Na ions | 0.74 |
| 2 kD FITC-PEG-COOH | 0.08 |
| 2 kD FITC-PEG-NH3 | 0.04 |
| 10 kD FITC- PEG | 0.02 |
| NADH | 0.09 |
| GFP fluorescent protein | 0.06 |

**Supplementary Table 1.** Ratio of solute concentration gradients in water (140 mM KCl) versus 10 gram% gelatin, on opposing sides of the interface between phases. This ratio approximates the ratio of diffusion coefficients in the two phases without compensation for solute binding by the gelatins. Estimated diffusion coefficients for free solutes are graphed in Fig. S10. In cases where experiments were performed both into and out of the gelatin phase, the values given are an average of the two ratios obtained.

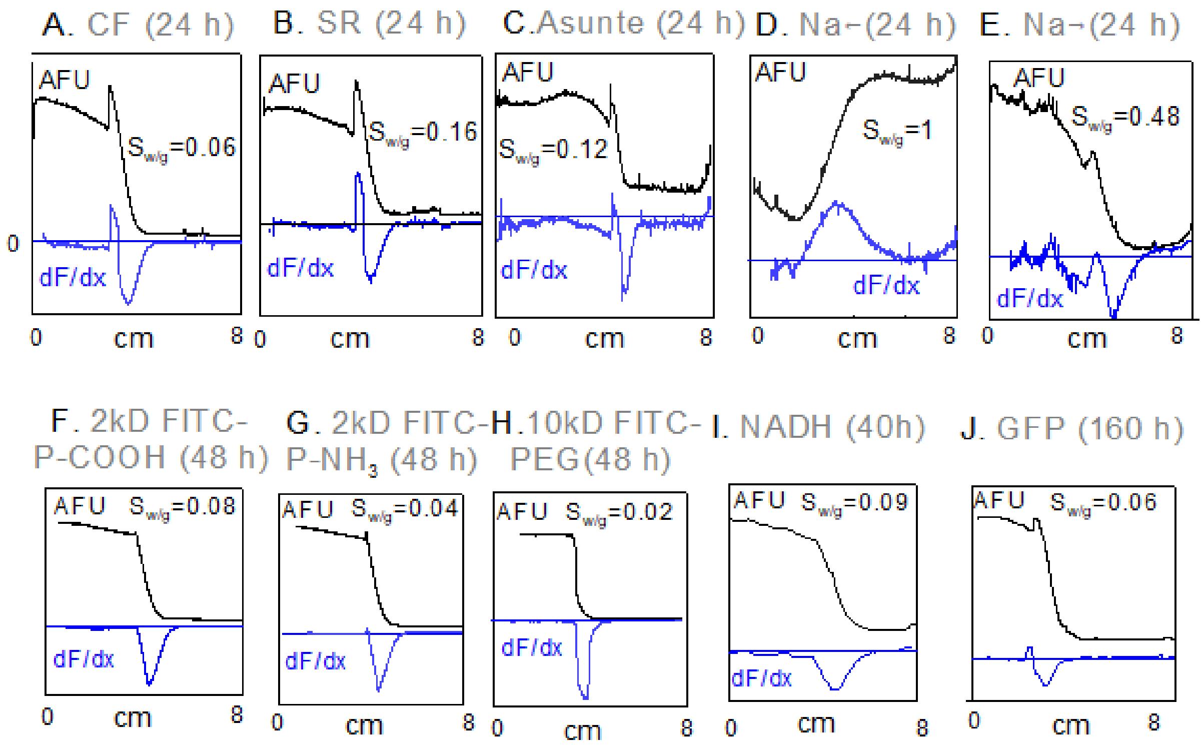


**Figure S6.** Diffusion of 8 fluorophores and Na ions in 8 cm glass pipettes in gelatins. All panels are oriented so that gelatin (10 gram%) is in the right half of the pipette and the left half is gelatin free. Fluorescence records are given in black, simulations are given as red lines, and the first derivative of fluorescence is given as a blue line. Solutions contained 140 mM KCl, 5 mM HEPES and 0.5 mM EGTA. **A-C.** Fluorescence profiles of three fluorophores 24 h after pipette loading: Carboxyfluorescein (CF,20 µM), sulforhodamine (SR, 20 µM), and ASG-2 (5 µM). $S_{w/g}$ values indicate that CF diffuses 17 times slower, SR 6.3 times slower and ASG2 8.3 times slower in gelatin than in gelatin-free solution. **D,E**. Profiles for Na diffusion after 24 h in pipettes containing ASG2 in both phases. Na (20 mM) was included in the gelatin phase in D and in the gelatin free phase in E. Gradients are 4.5 fold less steep than for dye in C, as expected for more rapid diffusion. ASG2 accumulates mildly (~10%) in the gelatin versus gelatin-free solution. Consistent with electrical measurement, Na diffusion is restricted by at most 50% in gelatin. **F-H**. Fluorescence profiles of 8 cm pipettes 48h after initiating diffusion of 2 kD FITC-PEG-COOH (20 µM), 2kD PEG-$NH_3$ (20 µM), and 10 kD PEG (20 µM). Accumulation in gelatin is very small or negligible, and ratios of fluorescence slopes are 0.08, 0.04 and 0.02, respectively, indicative of strong diffusion restriction in the gelatin phase. **I.** Diffusion profile of NADH (M.W. 663, 150 μM) 40 h after initiating diffusion from the gelatin-free phase into gelatin. NADH diffusion is restricted by 11 fold in the gelatin phase. **J.** GFP (27 kD) diffuses slowly but significantly into gelatin over 1 week. It is restricted in the gelatin versus gelatin free solution by 17 fold.

To estimate the diffusion coefficients of free solutes in these experiments, diffusion out of the gelatin phase was restricted by a variable fraction (F) that reconstructed as accurately as possible the rise of fluorescence at the interface to the gelatin. The diffusion coefficient in gelatin, used for the plot in Fig. S10, was then calculated as the diffusion coefficient in water times the slope ratio $S_{w/g}$ and F.

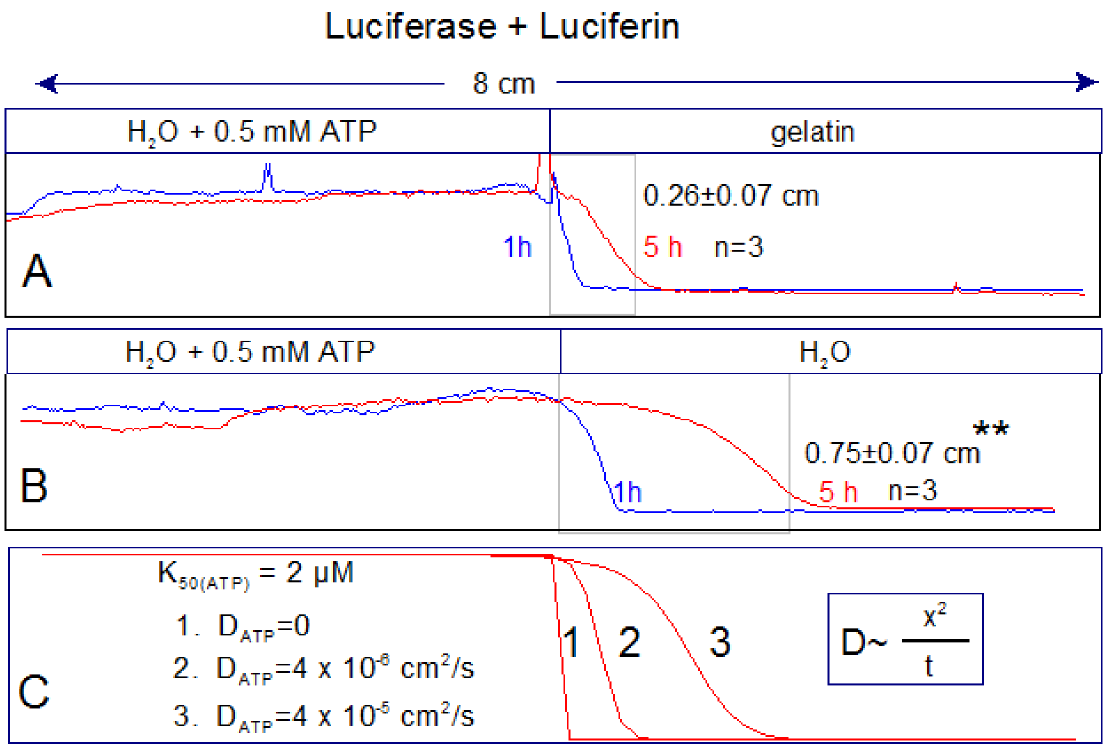


**Figure S7**. Optical measurement of ATP diffusion in 8 cm long pipettes, monitored via fire fly luciferase luminescence (Rasmussen and Nielsen, 1968). All solutions contained luciferase (150 ng/ml) and luciferin (15 µM). One group of pipettes (A) was loaded one-half with gelatin free solution containing 0.5 mM ATP and one-half with ATP-free gelatin. A second group (B) was loaded entirely without gelatin and with ATP in one-half of the pipette. Chemiluminescence was then evaluated after 1 and 5 h. The ATP gradient moved on average three-fold further in the water phase (B) than in the gelatin phase (A), indicative of a 9-fold smaller diffusion coefficient in 10 gram% gelatin. (C) Simulation of results with the diffusion coefficients indicated and with an ATP dissociation coefficient of 2 µM.

**Additional details.** Three pipettes were half-filled with gelatin and three pipettes were half-filled with gelatin-free solution. Then, solutions containing MgATP (0.5 mM), luciferase, and luciferin were carefully filled into the second halves of the tubes.. A complexity of this experiment is that the luciferase has high affinity for ATP, so that chemiluminescence has a non-linear dependence on ATP concentration. Nevertheless, as shown in Panels A and B of Fig. S7, the experiment demonstrates unambiguously that for a given amount of time, MgATP diffuses three times further in gelatin-free solution than in 10 gram% gelatin ($p<0.01$). Since the diffusion coefficient is proportional to the square of distance, the ATP coefficient in gelatin is 9 times smaller than in gelatin-free solution. We also verrifie this conclusion by simulation. Chemiluminescence was assumed to be proportional to ATP binding, and the $K_d$ for ATP was varied over a range of 1 µm to 20 µm. Regardless of the affinity, a threefold change of diffusion distance required that the diffusion coefficient was 9-times smaller in gelatin.

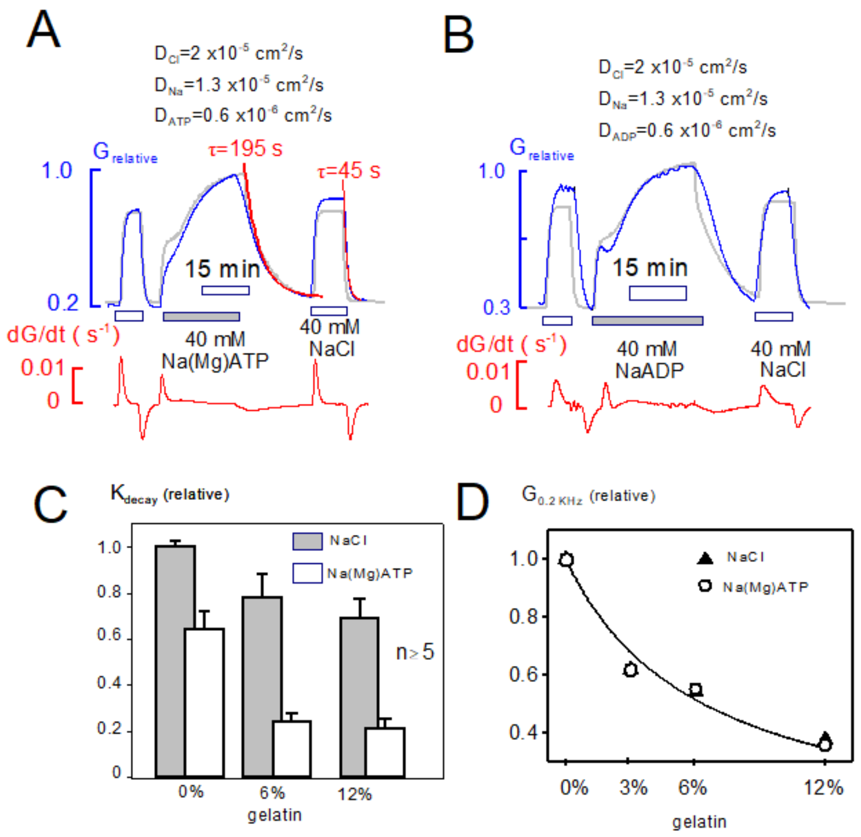


**Figure S8.** Comparison of NaCl, Na(Mg)ATP and NaADP diffusion in gelatin using conductance measurements to monitor diffusion. **A**. NaCl (40 mM) was applied and removed, followed by Na(Mg)ATP (40 mM) and then NaCl again. The gray curve is a simulation for a Na-anion pair in which the anion diffuses 10 times slower than Cl. Similar to the simulation, the conductance record for 40 mM Na(Mg)ATP rises in a biphasic fashion and declines >3-times slower than the records for NaCl. **B.** NaADP diffusion into a gelatin-filled pipette tip follows a similar pattern to Na(Mg)ATP. **C.** Comparison of NaCl and Na(Mg)ATP diffusion in pipette tips with 0, 6 and 12 gram% gelatin. For all results, n≥5. The decay constant for NaCl decreases by 25 and 41% in 6 and 12% gelatin, respectively, while the decay constant for Na(Mg)ATP is decreased by 80% in both gelatins. We mention that results for NADH were similar.

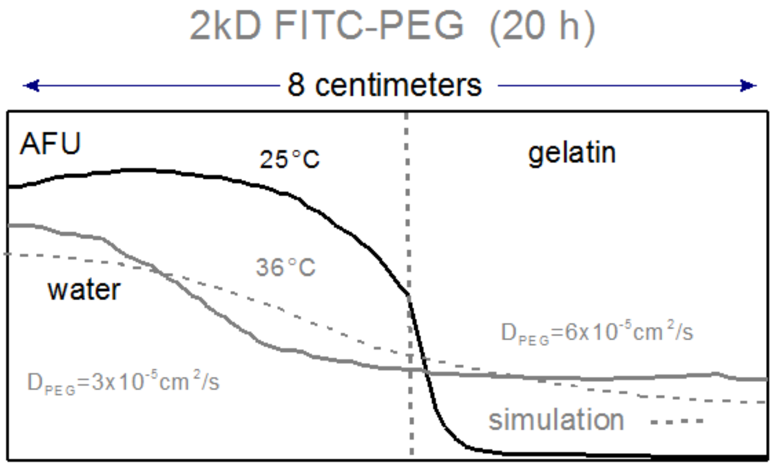


**Figure S9.** Diffusion restrictions in gelatin are lost by warming to 36°C, in close association with liquification of the gelatin. Diffusion of 2kD FITC-PEG at 25 and 36°C for 20 h. Six 8 cm pipettes (8 cm) were filled one half with 10 gram% gelatin and one half with gelatin-free solution containing 10 µM FITC-PEG. Three were incubated at 37°C and three at 25°C for 20 h. Gradients in pipettes incubated at 25 °C were very similar to those presented in Figs. S5 and S7 , diffusion being 5 times slower in the gelatin phase. In pipettes incubated at 37°C FITC-PEG diffused the entire length of the pipettes in 20h, so that fluorescence at the loaded end decreased by nearly 20% and increased by a similar amount at the unloaded end. To account for the data, the diffusion coefficient in the gelatin phase must be close to $10^{-4}$ $cm^2/s$. Possibly, therefore, solute movements are in some way facilitated in this condition. We conclude that diffusion restrictions in gelatin are dependent on gelatin cross-linkages and not on the molecular crowding per se of proteins in solution.

**Diffusion restrictions employing other macromolecules.** For brevity we describe only with words additional experiments in which we compared diffusion in gelatins with diffusion through several viscous macromolecular solutions. To do so, we employed pipettes prepared as in Figs. 1D and 1E, and we compared diffusion of Na(Mg)ATP and NMG-MES when the pipette tips were loaded with viscous solutions containing 10 gram% polyvinylpyrrolidone (360 kD), 5 gram% silica particles (200 nm, Aerosil), or 10 gram% fatty acid free albumin. Diffusion restrictions were small or even negligible, compared to diffusion in gelatins.

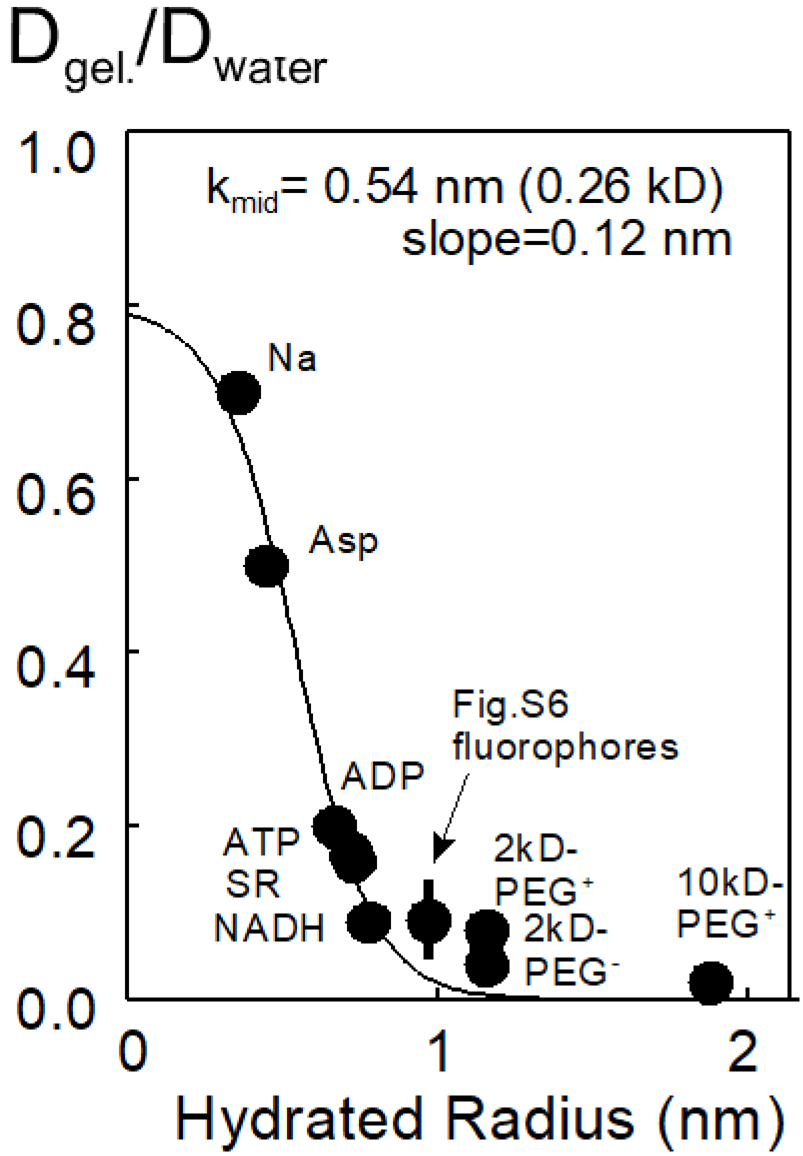


**Figure S10.** Ratio of estimated diffusion coefficients of selected compound in 105 gelatin versus free water (140 mM KCl). For Na, ATP, and NADH the estimated diffusion coefficients are an average of values obtained in both optical and electrical measurements.

**Reference List**


Iamshanova, O., P. Mariot, V. Lehen'kyi, and N. Prevarskaya. 2016. Comparison of fluorescence probes for intracellular sodium imaging in prostate cancer cell lines. *European biophysics journal : EBJ*. 45:765-777.

Rasmussen, H., and R. Nielsen. 1968. An improved analysis of adenosine triphosphate by the luciferase method. *Acta Chem Scand*. 22:1745-1756.

Robinson, R.A.S., R.H. 1965. Electrolyte solutions: The measurement and interpretation of conductance, chemical potential and diffusion in solutions of simple electrolytes. . Butterworths. 571 pp.

Swietach, P., K.W. Spitzer, and R.D. Vaughan-Jones. 2015. $Na^+$ ions as spatial intracellular messengers for co-ordinating $Ca^{2+}$ signals during pH heterogeneity in cardiomyocytes. *Cardiovascular research*. 105:171-181.

Wang, T.M., and D.W. Hilgemann. 2008. Ca-dependent nonsecretory vesicle fusion in a secretory cell. *J Gen Physiol*. 132:51-65.
